# Mechanistic Decoding of Pharmacodynamic Intratumoral Heterogeneity: A Pan-Cancer Regulatory Framework

**DOI:** 10.64898/2026.10.01.756124

**Authors:** Zhixiang Ren, Yiming Ren, Zeting Li, Shuang Ge, Yaxin Xu, Huan Xu

## Abstract

Intratumoral heterogeneity (ITH) drives therapy failure through divergent drug response propensities among coexisting subclones—a phenomenon we term pharmacodynamic intratumoral heterogeneity (PharmITH). While single-cell technologies have catalogued transcriptional heterogeneity, which molecular programs shape differential drug responses remains unresolved. Here we decode PharmITH by linking single-cell transcriptomes from 1,279 samples across 24 cancer types with drug-response profiles for 1,218 compounds. We identify 425 hub genes consistently associated with predicted treatment-response heterogeneity and with potential therapeutic relevance and perform multi-omics network analysis in lung adenocarcinoma (LUAD) integrating transcriptional regulation, somatic mutations, cell-cell communication (CCC), and alternative splicing. This analysis identifies four classes of candidate regulatory associations observed in the LUAD case study: 3 transcription factors establishing pre-existing drug-response programs, 41 genomic alterations amplifying pathway signaling through intermediate networks, 5 ligand-receptor pairs coupling microenvironmental cues to intracellular circuits, and 8 genes producing opposing isoforms via differential splicing. These findings converge on 6 core pathways, revealing that PharmITH arises from tumor-intrinsic mechanisms. Experimental validation and an online platform PharmITH for pan-cancer therapy enable state-resolved drug-response analysis and candidate-gene exploration. This approach transforms ITH from descriptive phenotyping to mechanistic dissection, providing actionable targets to overcome therapy resistance.

## 1. Introduction

ITH represents a fundamental barrier to effective cancer therapy, yet its impact extends beyond cellular diversity to encompass PharmITH—the coexistence of drug responder and drug non-responder subpopulations within individual tumors (1, 2). This pharmacological stratification means that while certain subpopulations respond to treatment, drug non-responder subpopulations survive and drive relapse (3–5). PharmITH thus transforms cancer from a disease treatable by single agents into an ecosystem requiring strategies that account for divergent drug-response propensities across coexisting populations. Effective precision oncology must therefore identify not merely which cellular states exist, but which molecular programs are consistently associated with differential drug responses.

Single-cell and spatial multi-omics technologies have mapped tumors at unprecedented resolution, revealing extensive transcriptional and phenotypic diversity (6, 7). However, these studies remain predominantly descriptive, cataloguing cellular states without systematically connecting state variation to predicted pharmacological differences (8). The critical gap lies in connecting molecular variation to pharmacological outcomes: among the myriad differences between subpopulations, which specific features are reproducibly associated with whether cells exhibit responder or non-responder states. Moreover, PharmITH involves regulatory features spanning genomic instability, transcriptional plasticity, epigenetic reprogramming, and microenvironmental interactions (9). Fragmented analyses examining individual molecular layers provide only partial views of the regulatory landscape surrounding drug-response heterogeneity, limiting candidate prioritization and subsequent functional investigation.

To address these limitations, we develop an integrated pipeline to characterize PharmITH by linking ITH-derived malignant-cell states to predicted drug-response phenotypes. We assemble a pan-cancer compendium of single-cell transcriptomes from 1,279 tumors spanning 24 cancer types, paired with drug response profiles for 1,218 compounds. By correlating gene expression variability across subpopulations with corresponding drug-response propensity patterns, we identify 425 hub genes that are consistently associated with predicted drug-response heterogeneity and may have potential therapeutic relevance. We further investigate the regulatory landscape surrounding these hub genes through four complementary layers—transcription factor regulation, somatic variation, CCC, and alternative splicing—to characterize the multi-omics features and pathway contexts associated with PharmITH.

In LUAD, this multi-omics analysis delineates four complementary regulatory dimensions surrounding PharmITH-associated hub genes: three recurrent transcription factors linked to state-specific drug-response programs; 41 variant events associated with differences in hub-gene activity and pathway states; five ligand–receptor pairs displaying differential communication patterns between predicted response groups; and eight genes exhibiting response-associated alternative-splicing patterns. Integration of these regulatory layers highlights six recurrent pathway categories. Wnt/β-catenin and PI3K/AKT/mTOR signaling are enriched in predicted responder-associated networks, whereas EMT, autophagy–lysosomal, NF-κB, and actin-cytoskeleton programs are enriched in predicted non-responder-associated networks. Together, these findings provide a multi-layer regulatory view of PharmITH and highlight candidate genes and pathways with potential functional and therapeutic significance.

## 2. Results

### 2.1. Pharmacological profiling of malignant subpopulations

To systematically dissect PharmITH, we develop a comprehensive analytical pipeline (**Figure. 1a**) that evaluates ITH across three key dimensions: functional phenotype, genotype and regulatory landscape. In phenotype layer, we integrate single-cell transcriptomic data from 24 cancer types sourced from Gene Expression Omnibus (GEO) (10, 11), with pharmacogenomic signatures of 1,218 compounds from the LINCS L1000 Connectivity Map database (12) to construct a pan-cancer atlas of PharmITH. As shown in **Figure. 1b**, this atlas incorporates 1,279 primary tumors and 5,297,049 cells, linking intratumoral transcriptional programs to in silico drug response variability.

**Figure. 1.**
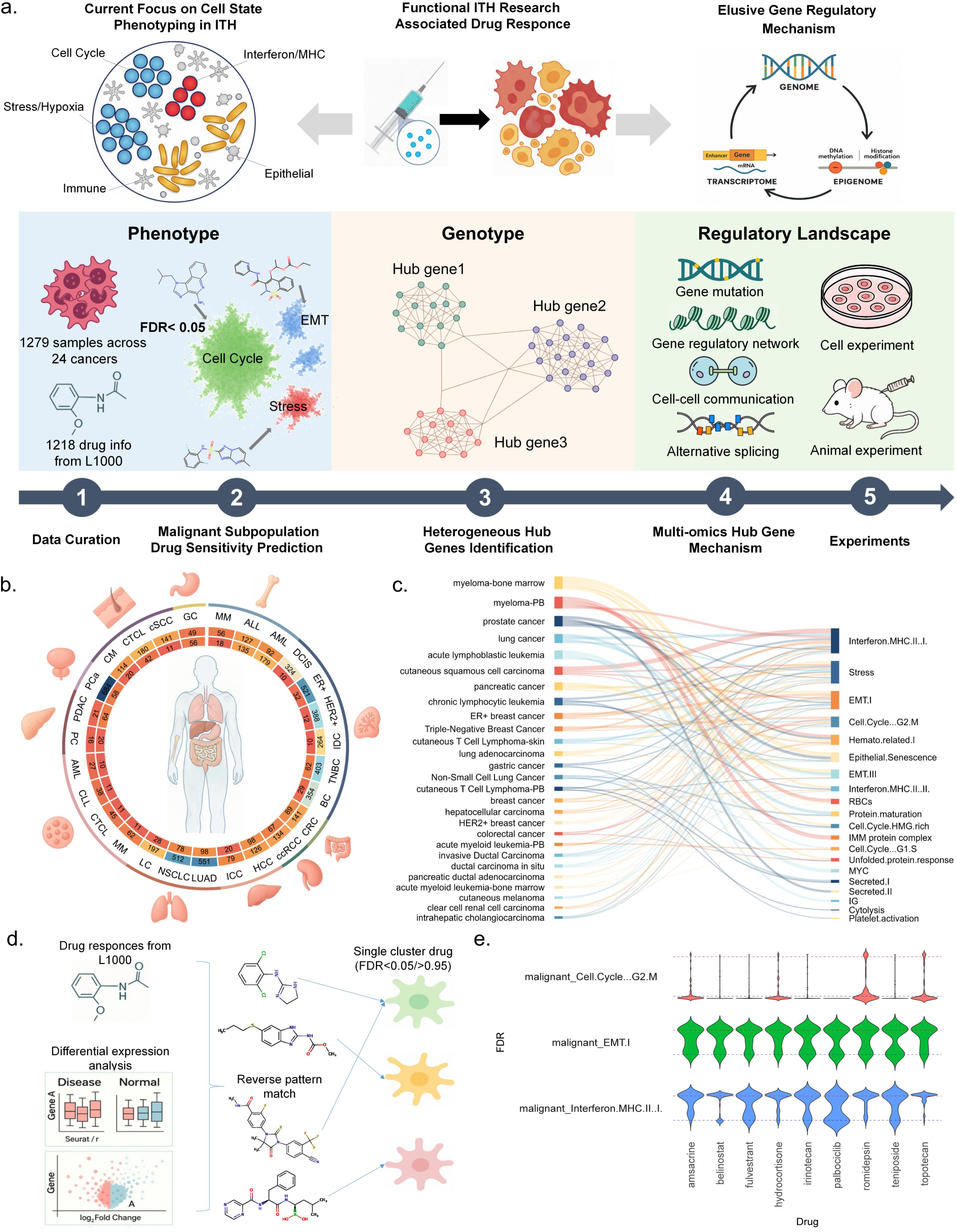
A pan-cancer regulatory framework for decoding pharmacodynamic intratumoral heterogeneity (PharmITH). (a) ITH is a fundamental cause of treatment failure. Although multi-omics integration has revealed phenotypic diversity in malignant cells, the molecular features associated with heterogeneous drug responses remain unclear. Here we establish a pan-cancer PharmITH analysis pipeline that innovatively integrates drug efficacy profiles with pan-cancer ITH phenotypes to: (1) predict subpopulation-specific drug-response propensity at the phenotypic level, (2) identify hub genes consistently associated with PharmITH at the genotypic level, and (3) characterize the regulatory landscape surrounding these hub genes across multiple molecular layers and associated signaling pathways. (b) The atlas integrates single-cell transcriptomic data from 1,279 primary tumors across 24 cancer types with pharmacogenomic profiles of 1,218 compounds from the LINCS L1000 database. The numbers in the outer ring indicate the number of drugs associated with each cancer type, whereas the numbers in the inner ring indicate the number of tumor samples included for the corresponding cancer type. Selected cancers are further subdivided by anatomical site. (c) 112 MPs, consensus expression programs shared across tumors, are annotated based on MP gene set and refined through GO pathway enrichment analysis. (d) Malignant cells are partitioned into subpopulations based on the defined MPs. Subpopulation-specific gene signatures are compared with L1000 drug perturbation profiles to compute connectivity scores, classifying drug responder status (FDR < 0.05) and drug non-responder status (FDR > 0.95) for each malignant subpopulation. (e) Pronounced differences in drug response are observed among malignant subpopulations exposed to the same therapeutic agent. In TNBC, Cell Cycle (G2/M) subpopulations show drug responder status for palbociclib, contrasting with drug non-responder status in EMT subpopulations.

Applying non-negative matrix factorization (NMF) to malignant cells from each tumor uncovers meta-programs (MPs) representing distinct cell states. We vary the number of NMF components (k=4–9) to capture heterogeneity at multiple resolutions, then consolidate similar gene programs across tumors into consensus MPs. **Figure. 1c** illustrates that 112 distinct MPs (**Supplementary Table 1**) is identified, each defined by a signature set of co-expressed genes. Gene set enrichment confirms that these MPs correspond to key oncogenic processes including cell-cycle proliferation, epithelial-mesenchymal transition (EMT) and inflammatory programs, consistent with known cancer cell state patterns (7).

We next assess drug response propensities of each malignant subpopulation defined by MPs. Adapting ASGARD (13) (**Figure. 1d**), we match each MP’s expression signature to L1000 drug perturbations to score efficacy for all 1,218 compounds per subpopulation. This distinguishes drug responder and drug non-responder subpopulations using strict false discovery rate (FDR) thresholds: drug non-responder status at FDR > 0.95 and drug responder status at FDR < 0.05. Notably, the prediction reveals pervasive intratumoral variation in drug-response propensity, even among subpopulations exposed to the same therapeutic agent. In TNBC samples, malignant cells from the proliferative G2/M subpopulation are classified as drug responders to cell-cycle inhibitors such as palbociclib, whereas coexisting cells from the EMT subpopulation are predicted to be drug non-responders to these same agents (**Figure. 1e**). These findings motivate deeper investigation into the molecular features associated with PharmITH.

### 2.2. Hub gene networks delineate divergent drug-response states

To identify molecular drivers of PharmITH, we first cluster drugs by mechanisms of action based on pathway-level changes in differentially expressed genes between drug responder and drug non-responder subpopulations (**Figure. 2, a, b, c**). Using these drug clusters, we perform meta-analyses on heterogeneity genes within each group to focus on shared biological processes. Compared with analysing all samples simultaneously, this approach enhances identification of convergent biological mechanisms underlying ITH. We construct protein-protein interaction networks from consistency genes and prioritize hub genes associated with PharmITH. Our pipeline identifies 425 hub genes across 24 cancer types (**Supplementary Table 2**). Literature mining reveals that 23% of these hub genes are known therapeutic targets and 36% are recognized biomarkers, a count that required validation in at least one of the 24 cancer types, supporting the utility of our pipeline in capturing PharmITH genes and underscoring the druggability of the identified hubs (**Figure. 2d, Supplementary Table 3**).

**Figure. 2.**
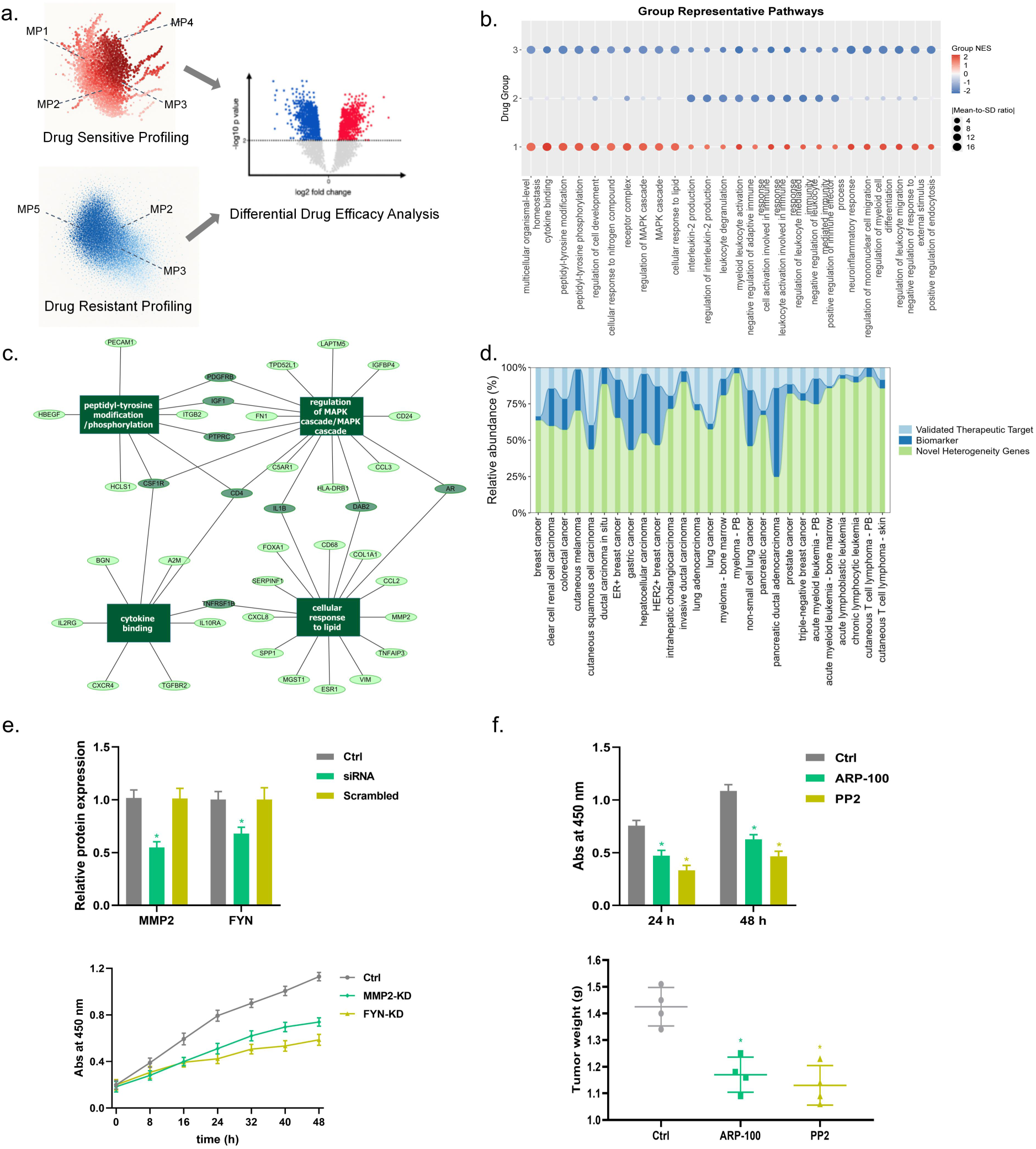
Hub genes delineate divergent drug responses. (a) Using TNBC as a representative example, differentially expressed genes between predicted responder and non-responder malignant subpopulations are integrated at the sample level to identify key genes associated with functional heterogeneity for specific drugs. (b) Heterogeneity genes for each drug are enriched in GO signaling pathways. The panel shows the pathway-enrichment profiles of three representative drug groups identified in TNBC samples. Each row represents a drug group, and each column represents an enriched GO pathway. Point color indicates the group-level normalized enrichment score (NES), whereas point size represents the absolute mean-to-standard-deviation ratio of the pathway NES values within the corresponding drug group. Hierarchical clustering based on normalized enrichment scores (NES) groups drugs with similar mechanisms, yielding 129 drug groups across 24 cancer types. (c) Heterogeneity genes showing consistency within the same drug group are subjected to functional enrichment analysis. The panel presents a representative GO pathway enrichment result of consistent heterogeneity genes identified in drug group 1 in TNBC. Dark-green gene nodes represent genes involved in multiple enriched pathways, whereas light-green gene nodes represent other consistent heterogeneity genes. Rectangular nodes represent significantly enriched biological processes. These genes were subsequently mapped to a PPI network for hub-gene prioritization. (d) Survey of PubMed-indexed studies (2018–2025) across 24 cancer types reveals that 23% of hub genes are therapeutic targets and 36% are biomarkers for prognosis, prediction, or diagnosis. Each stacked bar represents one cancer type and shows the relative proportions of hub genes classified as validated therapeutic targets, biomarkers, or novel heterogeneity-associated genes. (e) FYN and MMP2, which are hub genes in TNBC, are selected for experimental validation. Knockdown experiments in MDA-MB-231 TNBC cell lines using siRNA, compared with blank control and scrambled siRNA, demonstrate that silencing either FYN or MMP2 significantly suppresses malignant cell growth. The bar graph shows the semi-quantitative relative protein expression levels of MMP2 and FYN determined by ImageJ densitometric analysis of the Western blot bands. Original scans of western blot/gel images are shown in Figure S2. d. (f) Treatment with inhibitors targeting identified hub genes significantly suppresses tumor growth in both cell culture and xenograft models. Results are mean ± SD (n = 3 for in vitro experiments and n = 4 for animal studies). * p < 0.05 compared with Ctrl group.

To further validate our findings, we focus on two hub genes—FYN and MMP2—identified in TNBC analysis. FYN, a Src-family tyrosine kinase, has recently been identified as a mediator of drug tolerance and a potential combinatorial therapeutic target in TNBC (14, 15). Similarly, MMP2 is a matrix metalloproteinase that degrades extracellular matrix to facilitate tumor invasion, associates with increased metastasis. Although MMP2-associated pathways have been implicated in TNBC progression, direct validation of MMP2 itself as a therapeutic target in TNBC remains limited (16). We therefore carried out experimental validation to characterize the roles of FYN and MMP2 in TNBC.

As shown in **Figure. 2e**, knockdown of either MMP2 or FYN significantly suppresses tumor cell growth in the MDA-MB-231 TNBC cell line. ARP-100 was selected as a previously characterized MMP2 inhibitor (17, 18), whereas PP2 was selected as a Src-family kinase inhibitor with activity against Fyn (19–22). The docking models are provided as supplementary structural support for these established inhibitor–target relationships (Figure. S1). Both inhibitors significantly suppress tumor cell proliferation in vitro and reduce tumor progression in a murine subcutaneous TNBC model (**Figure. 2f and Figure. S2**). The uncropped and unprocessed original scans of the western blot images are provided in Figure S2. Collectively, these results provide proof-of-concept support for the functional relevance and pharmacological tractability of selected PharmITH-associated candidates in the tested TNBC model.

While hub genes represent key regulatory candidates in PharmITH, single-layer analyses yield fragmented insights. We therefore integrate transcriptional regulation, somatic variation, CCC, and AS to characterize the multi-layer regulatory landscape surrounding these hubs at the pathway level.

### 2.3. Intrinsic regulatory programs distinguish drug-response states

To characterize the regulatory landscape surrounding PharmITH-associated hub genes, we require paired scRNA-seq and scATAC-seq data for GRN inference. Five primary, treatment-naive LUAD samples in GSE274934 were selected as a representative cohort with paired scRNA-seq and scATAC-seq data for downstream regulatory annotation. All subsequent analyses across the four regulatory layers are conducted on this dataset. SCENIC+ (23) analysis reveals marked variation in enhancer accessibility between malignant subpopulations (**Figure. 3, a and b, Supplementary Note 1, and Figure. S3**), indicating that effective transcription factor inputs to hub genes differ by cell state. Such transcription factors orchestrate cell-intrinsic programs and represent potential therapeutic targets for reprogramming tumor cell states.

**Figure. 3.**
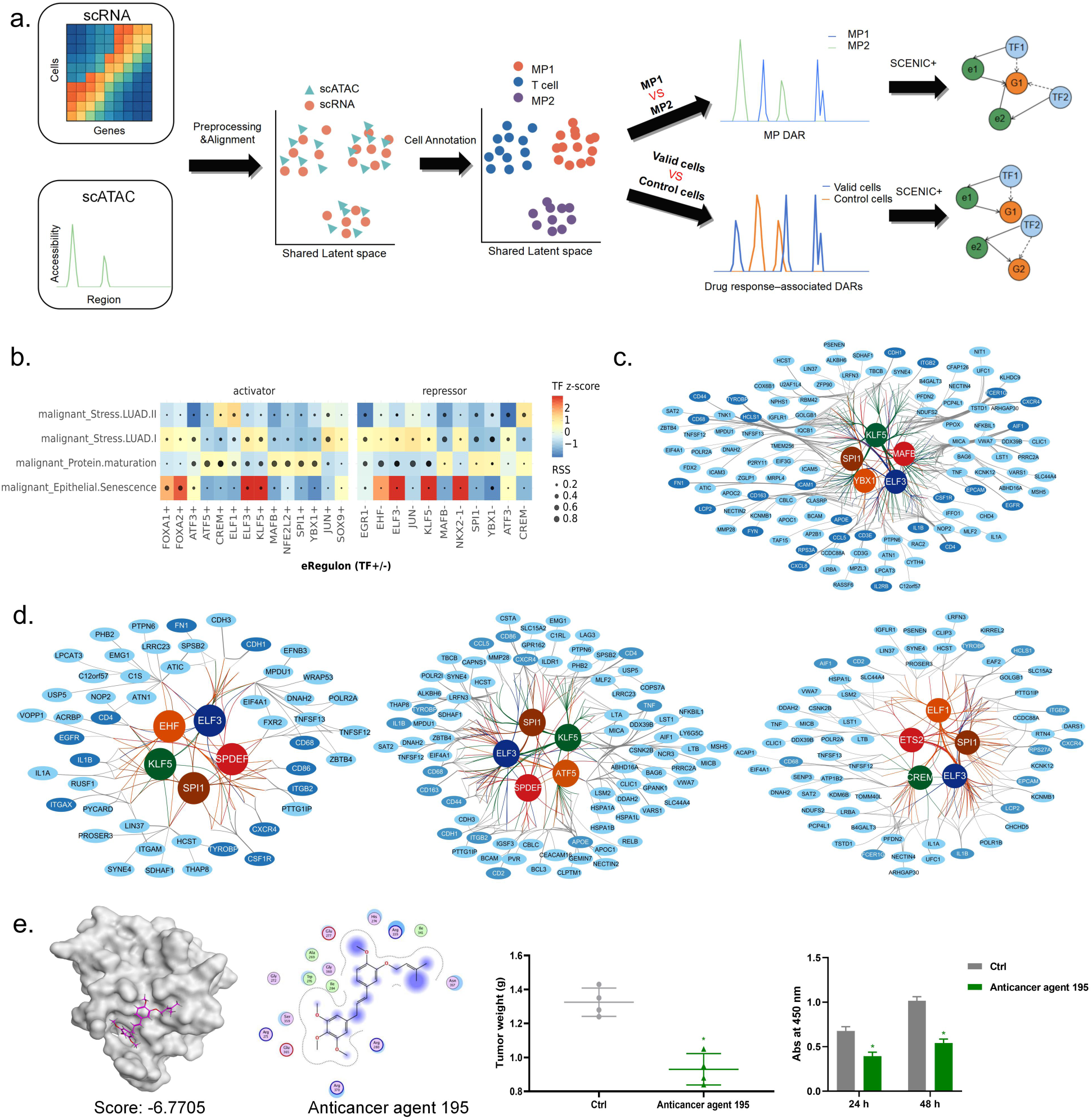
Intrinsic transcriptional and genetic programs distinguish drug response states. (a) Single cell ATAC sequencing (scATAC-seq) and single cell RNA sequencing (scRNA-seq) profiles are aligned in a shared latent space, with scATAC-seq cells annotated using the transcriptomic reference. CisTopic identifies differentially accessible regions among subpopulations, which are used by SCENIC+ to infer Transcription factor-Enhancer-Target gene triads linking transcription factor activity to hub genes. (b) The Regulatory Specificity Score (RSS) measures the specificity of hub gene regulation in each subpopulation. TF z-score represents normalized transcription factor expression levels within subpopulations. ‘+’ and ‘−’ indicate activating and repressive regulatory relationships between transcription factors and hub genes, respectively. The results of each drug group are shown in Figure. S8. (c) Confidence ranking of Transcription factor-Enhancer-Hub gene triads reveals that among the top 100 triads, five transcription factors—ELF3, YBX1, KLF5, SPI1 and MAFB—drive the highest number of hub gene regulatory events. Complete target gene sets for these transcription factors are shown, with hub genes highlighted in darker shading. (d) Analysis of the three most statistically significant drug groups (DG1, DG4 and DG5) demonstrates that ELF3, KLF5 and SPI1 associate with both subpopulation differentiation and PharmITH. (e) Treatment with the ELF3 inhibitor Anticancer agent 195 suppresses proliferation of LUAD cell lines in vitro and reduces tumor growth in a murine LUAD xenograft model in vivo. Results are mean ± SD (n = 3 for in vitro experiments and n = 4 for animal studies). * p < 0.05 compared with Ctrl group.

Using enhancer-centric network analysis, we identify five transcription factors—ELF3, KLF5, SPI1, MAFB and YBX1—as key regulatory anchors delineating malignant subpopulations (**Figure. 3c, Supplementary Note 2**). Strikingly, these transcription factors dominate ITH regulation and play leading roles in drug-response contexts. Across three drug groups (**Figure. 3d**), ELF3, KLF5 and SPI1 frequently emerge as recurrent regulators of drug-response efficacy (**Supplementary Note 3, Supplementary Table 4**). SPI1 (PU.1) functions as a master regulator of myeloid and immune-adhesion programs, driving expression of genes critical for immune signalling (24). ELF3, an epithelial-specific transcription factor, maintains epithelial lineage identity and suppresses EMT, thereby reinforcing drug responder epithelial states (25, 26). It also has previously been reported as a lineage-specific amplified oncogene and putative therapeutic target in LUAD (27). KLF5 acts as a context-dependent driver of epithelial proliferation and extracellular matrix interactions, contributing to the epithelial arm of drug response networks and modulating genes like CDH1 in coordination with other epithelial transcription factors (28). Together, these findings indicate that differences in cancer drug responsiveness are not random but instead orchestrated by specific transcription factor-driven programs pre-existing in certain malignant cell states.We validate ELF3 as a functional regulator of LUAD PharmITH. As shown in **Figure. S4**, knockdown of ELF3 can significantly suppress tumor cell growth in LUAD cell line. Moreover, the ELF3 inhibitor Anticancer agent 195, confirmed by molecular docking, significantly suppresses LUAD cell proliferation in vitro and reduces tumor progression in a murine subcutaneous LUAD model (**Figure. 3e**), demonstrating ELF3’s therapeutic potential. The original uncropped western blot images with molecular weight markers for ELF3 in A549 cells, as well as for MMP2 and FYN in MDA-MB-231 cells, are provided in **Figure S5.**

Somatic variants are additionally examined as a supplementary regulatory-context layer. We identify 41 variant events associated with three predicted drug-response groups. These variants generally occur outside the predefined hub genes and are associated with specific hub-gene activity and pathway states. Complete SComatic results and illustrative pathway annotations, including FAT1-associated YAP/TAZ activation and predicted verteporfin response (29–31), as well as TCIRG1-associated lysosome–autophagy programs and predicted responses to chloroquine or tamoxifen (32, 33), are provided in Supplementary Note 4 and Supplementary Table 5.

### 2.4. Intercellular and post-transcriptional regulation extends the PharmITH hub-gene landscape

Following the transcriptional and genetic analyses in Section 2.3, we next examine whether PharmITH-associated hub genes are embedded in additional regulatory contexts at the intercellular and post-transcriptional levels. Cell-cell communication and alternative splicing therefore constitute the third and fourth layers of our hub-gene-centered regulatory landscape, complementing transcription factor regulation and somatic variation.

Ligand-receptor analysis of hub genes reveals divergent cell-cell communication across four malignant LUAD subpopulations (**Figure. 4, a and b**). The Epithelial.Senescence subpopulation exhibits a senescence-associated secretory phenotype characterized by elevated JAK-STAT cytokine signalling and frequent contacts with CD8+ T cells, NK cells and myeloid cells. These senescent-like tumor cells secrete inflammatory mediators recruiting immune effectors, consistent with immunosurveillance, yet retain active EGFR signalling despite growth arrest. In contrast, the Protein.maturation subpopulation primarily engages monocytes and fibroblasts, featuring strong CD44-extracellular matrix (ECM) interactions. Collagen ligands from perivascular stromal cells binding CD44 suggest a mesenchymal, matrix-attached phenotype providing pro-survival signals and promoting immune evasion. The two stress-adapted subpopulations (Stress I and II) engage in paracrine signalling, secreting high levels of CXCL8 (IL-8) and CCL5 that bind endothelial ACKR1. IL-8 recruits neutrophils, dampening cytotoxic immunity and promoting EMT. These stress-state cells upregulate NF-κB/TNFα inflammatory pathways and interact with immunosuppressive tumor-associated macrophages (**Supplementary Note 5**).

**Figure. 4.**
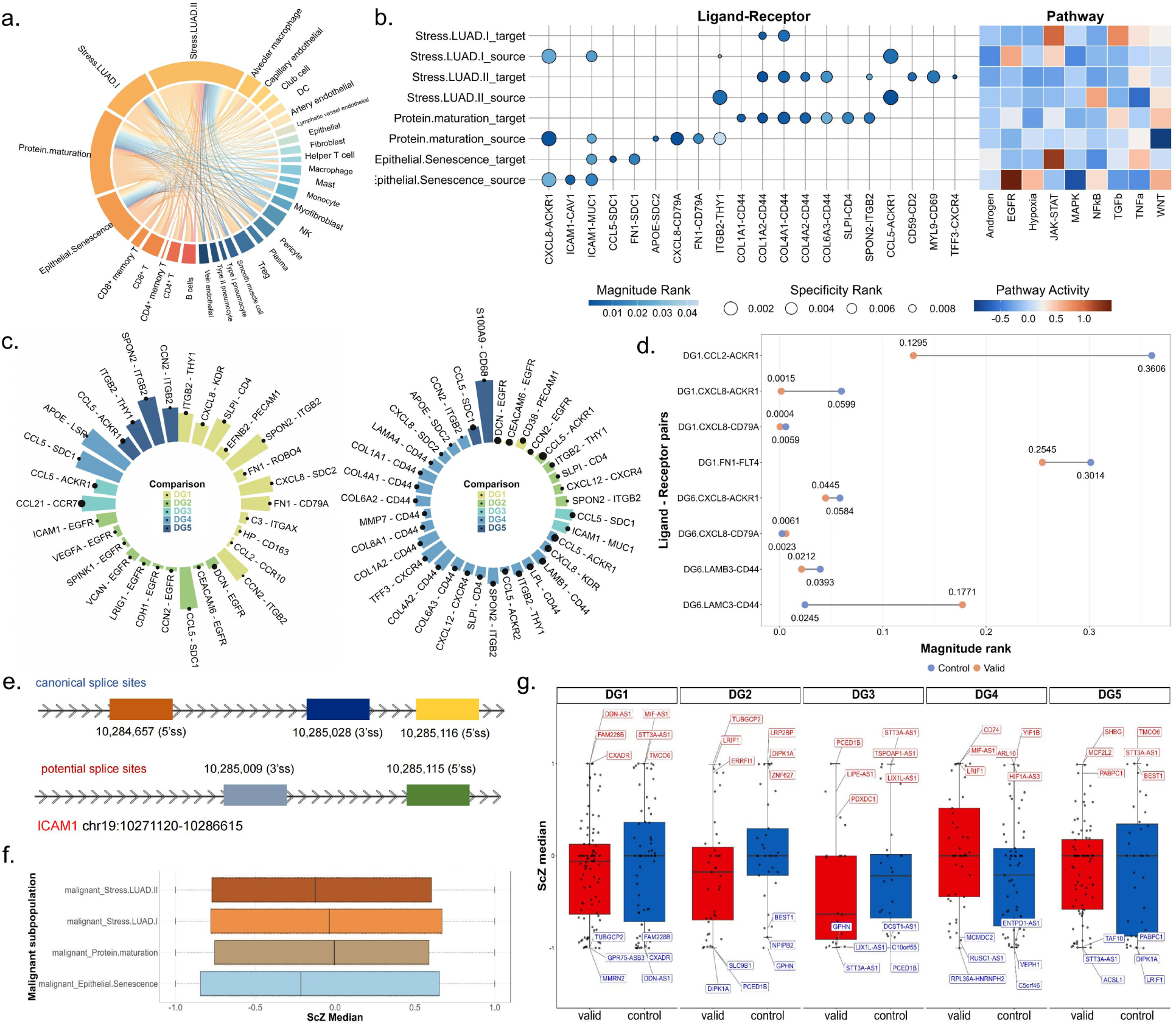
Extrinsic signaling and post-transcriptional rewiring diversify hub gene functions. (a) Differential signaling interactions across malignant subpopulations. Four malignant subpopulations exhibit distinct signaling interaction profiles with microenvironmental cell types. (b) Top-ranked pathways and ligand-receptor pairs in which malignant subpopulations act as signal senders or receivers. (c) Left panel shows unique, highly significant ligand-receptor pairs in drug responder subpopulations; right panel shows those in drug non-responder subpopulations. Notably, FN1-CD79A (drug responder group), ICAM1-MUC1 and TFF3-CXCR4 (drug non-responder group) also function as hub factors. (d) The ITH-driving ligand-receptor pairs CXCL8-CD79A and CXCL8-ACKR1 display opposite interaction patterns between drug responder and drug non-responder groups. (e) Differential splicing patterns of hub gene ICAM1 between drug responder and drug non-responder subpopulations. (f) The scz median value quantifies the deviation of a cell’s splicing pattern from the population average. In all four malignant subpopulations, there is a preferential shift towards short-intron splicing events. (g) Compared with the drug responder group, drug non-responder subpopulations show increased propensity for long-intron splicing events. This indicates that splicing shifts rewire gene function, modulating pathways such as EMT and survival signaling.

Examining cell-cell communication in relation to therapy response reveals overlap between ITH-associated and PharmITH-associated interactions (Supplementary Table 6). Among hub gene-mediated ligand-receptor pairs, five showed distinct patterns between drug responder and drug non-responder subpopulations (Supplementary Note 6). Drug responder subpopulations uniquely display FN1-CD79A (34), while drug non-responder subpopulations harbor ICAM1-MUC1 and TFF3-CXCR4 (Figure. 4c). CXCL8-ACKR1 shows a marked response-associated shift in interaction strength (Figure. 4d). These results place PharmITH-associated hub genes within distinct intercellular communication programs, extending the hub-gene regulatory landscape beyond the cell-intrinsic transcriptional and genetic layers characterized above.

We next asked whether the same drug-response states are also distinguished at the post-transcriptional level. Alternative splicing analysis reveals another layer of ITH. All four subpopulations favour shorter intron splicing (median splicing z-scores < 0; **Figure. 4f**), yet splicing programs diverge between drug responder and drug non-responder tumors (**Supplementary Table 7**). Drug responder cells favour longer intron-retained isoforms, whereas drug non-responder cells produce shorter variants conferring survival advantages (**Figure. 4g**). ICAM1 undergoes multiple splicing changes at five sites (three canonical, two novel) and serves as a communication hub bridging tumor cells and endothelium in the senescent cluster (**Figure. 4e**), exemplifying how hub genes operate across layers through intercellular signalling and alternative splicing modulation. Moreover, some genes exhibit completely opposite splicing patterns between the drug non-responder and drug responder subpopulations (**Supplementary Note 7**). For instance, CXADR shows opposite splicing patterns: drug responder cells predominantly express a full-length tight-junction adhesion receptor isoform, whereas drug non-responder cells favour a shorter variant lacking the complete CAR domain, consistent with differences in EMT and TGF-β signaling. Similarly, the long non-coding RNA STT3A-AS1 generates a longer intron-retained isoform in drug non-responder cells, linking its splicing pattern to STT3A-related glycosylation programs (35). Together, these isoform shifts reveal a post-transcriptional regulatory layer that may contribute to divergent drug-response phenotypes.

Together with the transcription factor and somatic-variation analyses, these intercellular and post-transcriptional results complete four complementary regulatory dimensions surrounding PharmITH-associated hub genes and provide the multi-layer evidence integrated at the pathway level.

### 2.5. Multi-layer regulatory signals converge on core PharmITH pathways

Previous analyses identified regulatory events across four molecular layers. To determine whether these distinct regulatory features converge at the functional level, we independently mapped transcription factor regulation, somatic variation, cell-cell communication, and alternative splicing to canonical signaling pathways and summarized recurrent pathway-level patterns across predicted response states. Integrated analysis reveals that multi-layer PharmITH-associated signals converge on a limited set of core signaling pathways (**Figure. 5a**). Drug responder subpopulations are significantly enriched for Wnt/β-catenin and PI3K/AKT/mTOR signaling, whereas drug non-responder subpopulations are primarily characterized by EMT, autophagy-lysosomal, NF-κB, and actin-cytoskeleton programs (**Supplementary Note 8**).

**Figure. 5.**
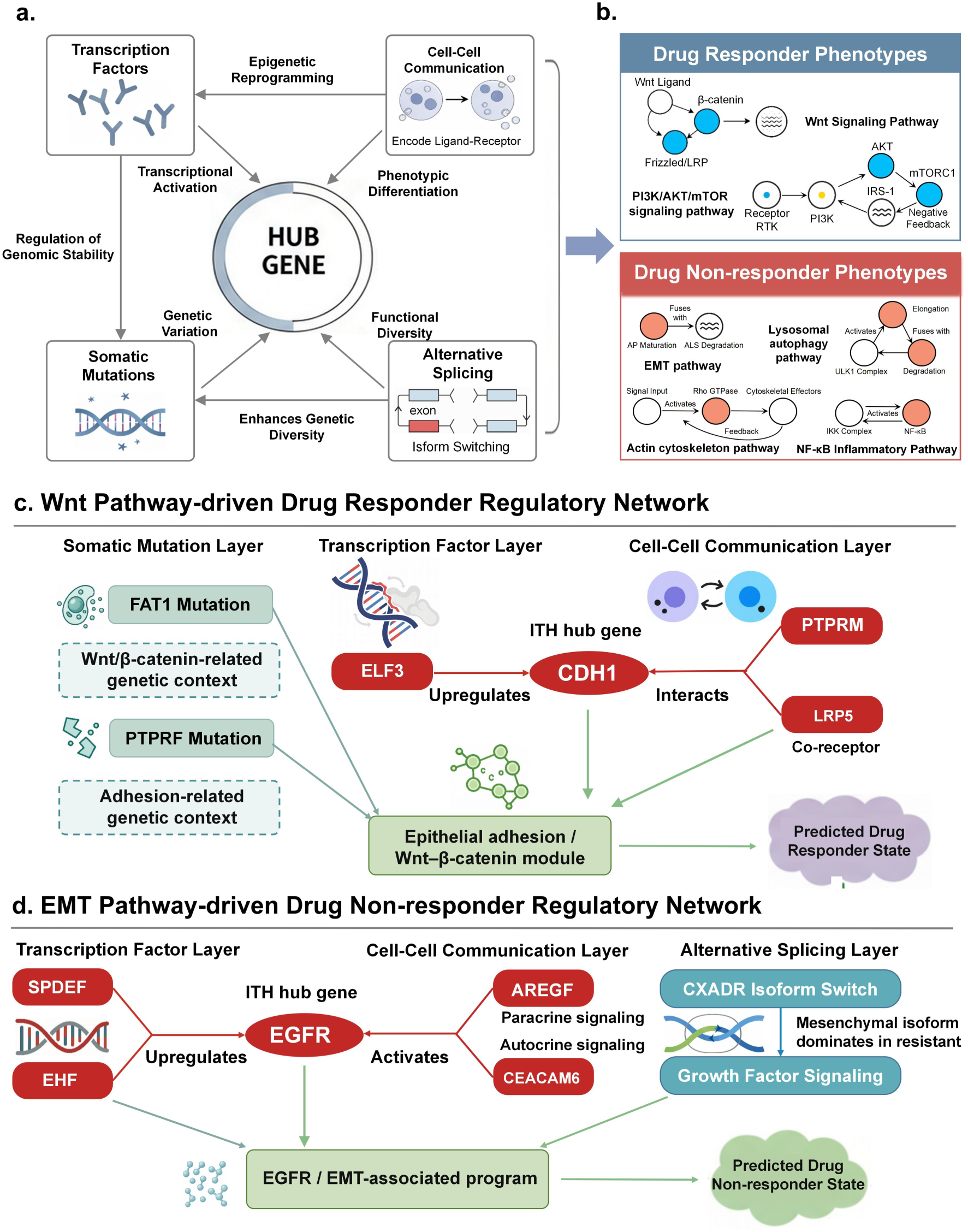
Multi-layer regulatory signals converge on core pathways shaping the PharmITH landscape. (a) Integration of PharmITH-associated regulatory features across four molecular layers—transcription factor regulation, cell-cell communication, alternative splicing, and somatic variation—to identify recurrent pathway-level convergence. (b) Drug responder phenotypes are enriched in Wnt/β-catenin and PI3K/AKT/mTOR pathways, whereas drug non-responder phenotypes are characterized by EMT, lysosomal autophagy, NF-κB, and actin-cytoskeleton remodeling pathways. (c) Representative Wnt/β-catenin-centered regulatory module associated with the drug responder phenotype. ELF3 is linked to CDH1 in the transcription factor layer; PTPRM-and LRP5-associated interactions connect cell adhesion with Wnt-related signaling in the cell-cell communication layer; and FAT1 and PTPRF variants provide additional genetic features within this regulatory context. These features converge on an epithelial adhesion/Wnt–β-catenin program associated with the predicted responder state. (d) Representative EGFR/EMT-centered regulatory module associated with the drug non-responder phenotype. EGFR is linked to SPDEF and EHF in the transcriptional layer, while AREG-EGFR and CEACAM6-EGFR interactions are identified in the communication layer. At the post-transcriptional level, CXADR displays response-associated alternative splicing linked to epithelial-mesenchymal plasticity. These features converge on an EGFR/EMT program associated with the predicted non-responder state.

For example, in drug responder LUAD subpopulations, multiple regulatory layers converge on an epithelial adhesion/Wnt–β-catenin module centered on CDH1 (**Figure. 5b**). The epithelial transcription factor ELF3 positively regulates CDH1 expression, reinforcing epithelial phenotype. At the cell-cell communication layer, PTPRM- and LRP5-associated interactions connect cell-adhesion features with Wnt-related signaling. At the genetic layer, FAT1 and PTPRF variants provide additional variation within this adhesion-associated regulatory context; notably, FAT1 has an established role in β-catenin/Wnt regulation (36). Together, these independently identified transcriptional, intercellular, and genetic features converge on a Wnt/β-catenin-centered regulatory module associated with the predicted drug responder state.

A complementary pattern is observed in drug non-responder LUAD subpopulations, where multi-layer signals converge on an EGFR/EMT-centered regulatory module (**Figure. 5c**). Drug non-responder cells show high expression of the EGFR-associated regulators SPDEF and EHF. Cell-cell communication analysis identifies paracrine AREG-EGFR signaling between malignant and endothelial cells and CEACAM6-EGFR interactions within malignant populations. At the post-transcriptional level, CXADR exhibits response-associated alternative splicing, consistent with its established involvement in epithelial-mesenchymal plasticity (37). Together, these transcriptional, intercellular, and post-transcriptional features converge on an EGFR/EMT-associated program characteristic of the predicted drug non-responder state.

Beyond these representative networks, the other recurrent pathways are similarly supported by regulatory features from multiple analytical layers. PI3K/AKT/mTOR signaling forms an additional responder-associated module, whereas autophagy-lysosomal, NF-κB inflammatory, and actin-cytoskeleton programs recurrently characterize non-responder-associated networks **(Supplementary Note 8)**. By integrating the four molecular layers, we reveal that PharmITH is organized around coherent functional modules spanning genomic, transcriptional, communicative, and post-transcriptional dimensions. Hub genes serve as integrative anchors through which diverse molecular features converge on these core pathway programs, highlighting regulatory contexts with potential functional and therapeutic significance.

### 2.6. Pan-Cancer Computational Platform for Precision Oncology

We present a modular computational platform PharmITH for precision oncology that integrates pan-cancer multi-omics analysis with PharmITH profiling. The platform, as illustrated in Figure. 6, comprises three core layers: (1) a Data Management Layer housing a multi-omics data warehouse (scRNA-seq, scATAC-seq), drug efficacy profiles encompassing small molecules and cell line perturbations, and a curated knowledge base; (2) a Mechanism Analysis Layer that resolves cellular identity, delineates malignant subpopulations, and performs multi-dimensional regulatory inference including somatic mutation detection, transcription factor networks, CCC, and AS; and (3) downstream translational applications for precision drug targeting within malignant subpopulations and therapeutic target discovery.

**Figure. 6.**
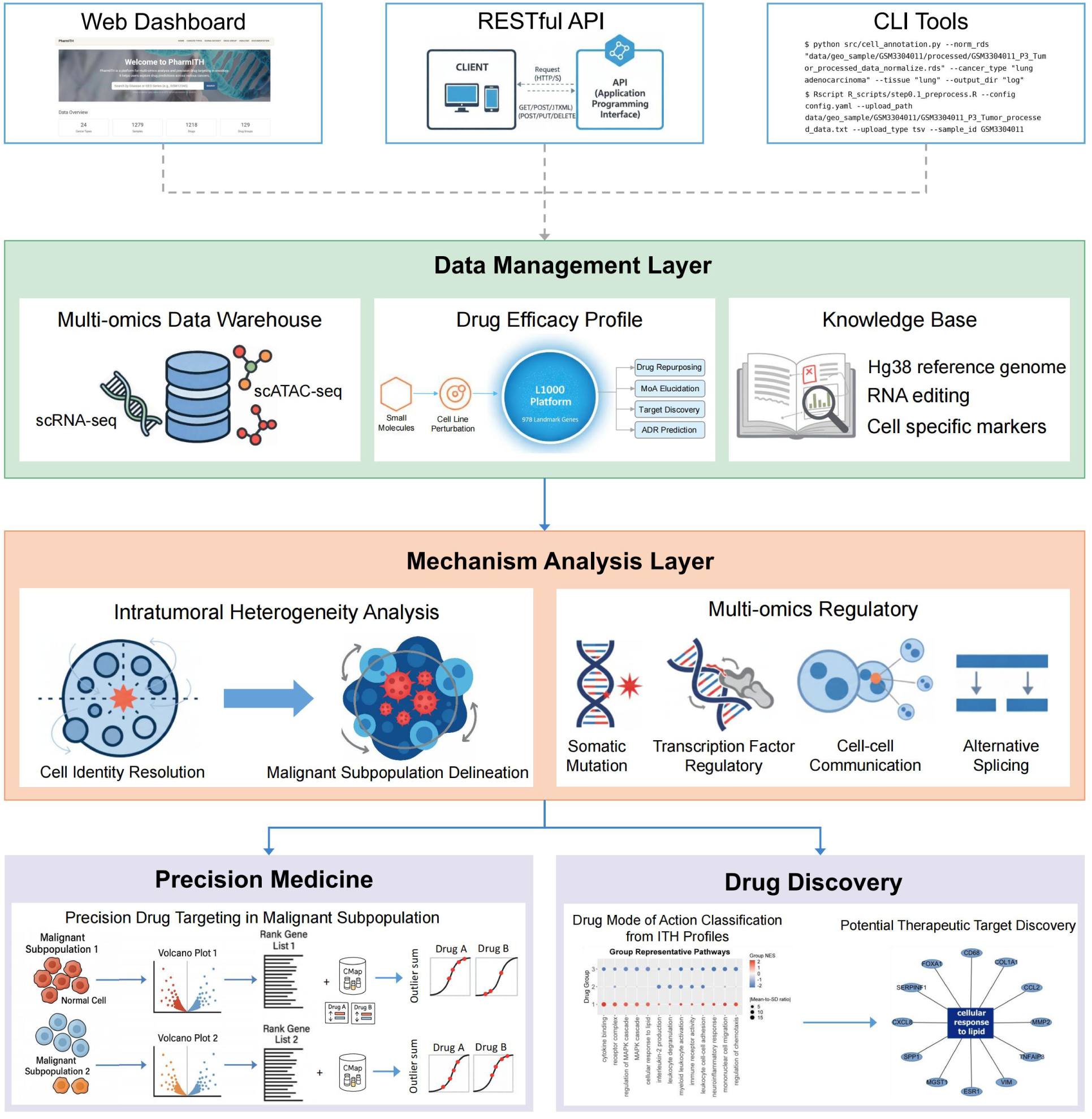
Pan-Cancer Computational Platform for Precision Oncology. The PharmITH platform comprises three computational layers supporting mechanism-driven precision oncology. The Data Management Layer integrates multi-omics data (scRNA-seq, scATAC-seq), drug efficacy profiles (small molecules, cell line perturbations) and a curated knowledge base. The Mechanism Analysis Layer resolves cellular identity, delineates malignant subpopulations through ITH profiling and performs multi-dimensional regulatory inference across somatic mutations, transcription factor networks, cell-cell communication and alternative splicing events. Two principal translational applications are enabled: (1) precision drug targeting within malignant subpopulations through correlation of ITH-derived MPs with drug efficacy data, and (2) mechanism-driven therapeutic target discovery through integrated regulatory network analysis. Three access modalities—web dashboard, RESTful API and command-line interface—support both interactive exploration and programmatic workflows for scalable pan-cancer analyses.

Architecturally, the platform employs a service-oriented design with FastAPI-based RESTful endpoints, Python core analytics modules, and R interoperability via subprocess bridges under a unified JSON/CSV schema. Three access modalities—web dashboard, RESTful API, and command-line interface—decouple computation from visualization, enabling both interactive exploration and programmatic workflows. This design supports scalable cohort-level analyses while maintaining extensibility for emerging single-cell technologies.

The platform supports two principal translational applications: precision drug targeting and mechanism-driven target discovery. For precision drug targeting, tumor scRNA-seq data are used as input, and the platform outputs malignant MP states, predicted drug-response propensities, candidate drugs, and corresponding visualizations. By correlating ITH-derived MPs with drug efficacy profiles, it identifies subpopulation-specific therapeutic vulnerabilities and enables precision drug assignment tailored to malignant cell states. For target discovery, cancer–drug-group results or predefined drug groups are used as input, and the platform outputs PharmITH-associated hub candidates, associated pathways, and supporting evidence layers. Through integrated regulatory network analysis, it prioritizes candidate genes associated with PharmITH, supporting drug repurposing and mechanism-of-action studies. The web interface and principal functions are available at pharmith.net, and complete implementation details and data can be obtained from the corresponding author.

## 3. Discussion

ITH is widely recognized as a major driver of therapy failure and cancer progression. Here, we extend descriptive ITH analysis by integrating single-cell transcriptomes with pharmacogenomic perturbation data to prioritize gene programs and pathways associated with divergent predicted drug-response propensities among coexisting malignant subpopulations. Gavish et al. characterized recurrent transcriptional programs underlying pan-cancer ITH, whereas ASGARD and DrInsight primarily use disease–drug expression reversal to prioritize candidate compounds (7, 13, 58). PharmITH adds a within-tumor, state-resolved comparison under the same drug reference, followed by cross-drug consistency and PPI-based candidate prioritization. Notably, malignant cell subpopulations are stratified into over a hundred MPs, revealing distinct cell-state networks linked to drug responder or drug non-responder status for particular compounds. The key hub genes emerging from these analyses represent prioritized candidate nodes and include kinases, proteases, and transcriptional regulators. Thus, beyond observing ITH, we prioritize candidate genes and pathways associated with predicted drug-response heterogeneity. Literature annotation and experimental evaluation support the further investigation of selected candidates as potential therapeutic targets.

However, the interpretation of these state-resolved response differences depends on the perturbational reference used. L1000 signatures are mainly derived from cancer cell lines under fixed-dose and short-term treatment conditions, with landmark genes directly measured and the remaining transcripts inferred. They therefore do not fully capture dose–response relationships, pharmacokinetics, metabolism, tissue distribution, immune or stromal effects, or long-term clinical resistance. Nevertheless, the present study focuses on tumor-cell-intrinsic states, which are compatible with the cancer-cell-line context of L1000, while its relatively standardized experimental conditions provide a unified pharmacological reference coordinate for comparing coexisting malignant states within the same tumor. In future studies, the perturbation input of PharmITH can be replaced with directional drug-response signatures derived from patient-derived organoids, PDX models, or clinical pre- and post-treatment samples.

While our study provides comprehensive insights into PharmITH, some challenges remain to be addressed. The lack of matched multi-omics data such as proteomic, mutational, or spatial data limits our ability to explore mechanisms comprehensively, and LUAD was therefore used as a representative multi-omics case study. The resulting regulatory associations require replication in larger independent cohorts and additional cancer types as suitable paired datasets become available. Beyond this cohort-level validation, the present multi-layer analyses identify pathway-level convergence but do not establish complete causal chains among regulatory events. Future studies should therefore combine subpopulation-specific perturbation experiments and time-series analyses with validation in independent cohorts to determine the temporal and causal relationships linking these regulatory layers to PharmITH. The core workflow also relies on CopyKAT-based malignant-cell identification. CopyKAT may have reduced discriminatory power in tumors with low CNV burden, very high tumor purity, or no adequate normal reference; future analyses should combine CNV inference with tumor markers and genomic or pathological validation. Moreover, an unexpected finding is that many hub genes are immune-related, even though the MP signatures are defined to minimize immune contributions. Further examination showed that these genes are concentrated in chemokine, cytokine, MHC/antigen-presentation, inflammatory, and adhesion programs expressed by malignant cells themselves (38). Immune-related proteins also tend to have greater connectivity in protein–protein interaction databases, which may slightly amplify their weight during hub-node ranking. The combination of malignant-cell immune-related expression programs and PPI connectivity bias may therefore explain this enrichment. Future studies will extend the framework to immune and stromal cells when suitable cell-type-specific perturbation data become available.

## 4. Methods

### 4.1. Cancer and drug perturbation data curation

To achieve results that closely recapitulate human cancer responses, human scRNA-seq datasets associated with cancer are obtained from the GEO (10) through March 12, 2025. Exclusions comprise data from cell lines, organoids, sorted cells, spatial transcriptomics, nuclear RNA, metastatic cancers, or treatment interventions. Samples with fewer than 200 cells are also discarded to ensure analytical robustness. To compute drug–gene connectivity scores based on disease and drug-perturbed expression profiles, we apply additional filtering to retain only samples with matched normal tissue scRNA-seq data and corresponding drug perturbation information from the LINCS L1000 (12). This curation process yields 127 projects spanning 24 distinct cancer types, encompassing 1,279 samples and a total of 5,297,049 cells. Additionlaly, we predict therapeutic scores for 1,593 FDA-approved drugs across various cancers using L1000 project data. A total of 1,218 drugs demonstrating clear tumor non-responder (FDR > 0.95) and tumor responder (FDR < 0.05) classifications are retained for subsequent analysis.

### 4.2. Quality control and normalization

We employ the Seurat package (v5.2.1) for quality control and normalization. Stringent cell-level filters are applied to exclude low-quality cells and multiplets. Specifically, cells with fewer than 200 or more than 5,000 detected genes (nFeatures) are removed, and cells containing fewer than 500 total RNA counts are filtered out. Gene-level quality control retains only genes detected in at least three cells. Normalization is then performed using the standard LogNormalize method as implemented in Seurat.

### 4.3. Cell type annotation

Following previous pan-cancer applications (39–41), malignant cells are identified using the R package CopyKat (v1.1.0) (42) to infer copy number variation (CNV) patterns within individual cells. The copykat function is employed with the following parameters: id.type = “S”, ngene.chr = 5, win.size = 25, KS.cut = 0.1, and distance = “euclidean”. Only cells classified as aneuploid were retained as malignant, whereas diploid or undefined cells were excluded from the main malignant-cell analysis. Non-malignant cells are annotated using the pre-trained deep neural network model scBERT (43), which operates independently of curated marker genes and effectively mitigates batch effects. To ensure tissue specificity across diverse cancer types, we collect annotated data from 11 organs (bone marrow (44), breast (45), colon (46), kidney (47), liver (48), lung (49), PB (44), prostate (50), stomach (51), pancreas (50) and skin (50)). To minimize technical variations, only primary data processed with the same sequencing platform, reagent kit version, and Cell Ranger version are retained. The base model, pre-trained on PanglaoDB, is fine-tuned for each organ to optimize performance for specific data characteristics and cell types, resulting in annotation of 64 cell types (**Figure. S6**). Annotation accuracy is further enhanced by integrating cell-type-specific markers from Cell Taxonomy (52), CellSTAR (53), and singleCellBase (54). Cells lacking expression of corresponding markers are labeled as “unassigned.”

### 4.4. Identification and characterization of malignant MPs

Malignant cells across 24 cancer types are profiled to identify MPs using GeneNMF (55). GeneNMF extends the consensus NMF approach and, unlike the method described by Gavish et al. (7), enables simultaneous clustering of programs across multiple k-values without pre-screening for robustness, thereby achieving greater specificity. Specifically, the 2,000 most variable genes are selected, and NMF is performed with k-values ranging from 4 to 9, yielding 39 initial programs per sample. These programs are then clustered based on shared genes to define refined MPs. Clustering parameters including the number of MPs, similarity metrics between clusters, and the extent of MP signature overlap are optimized using grid search. The Calinski-Harabasz index is employed to assess clustering solution quality. To ensure robust MP definitions, we exclude those with fewer than five specific genes, sample coverage below 30%, or a negative average silhouette coefficient. MP activity within individual cells is quantified using UCell-derived MP signature scores. Genes exhibiting significant differential expression between the top and bottom 20% of cells based on these scores are designated as MP-specific genes. Functional characterization of MPs is performed using an enrichment analysis approach inspired by 3CA (7). Gene sets derived from the 3CA-predefined MPs (7) are used for enrichment analysis, and the cell subpopulations most significantly enriched for each MP are assigned the corresponding functional label. Finally, following the strategy outlined by Jiang et al. (56), MP-specific genes serve as input for AddModuleScore to categorize malignant cells into distinct functional states.

### 4.5. Malignant subpopulation drug-response classification

To classify drug response in malignant subpopulations, we first perform differential gene expression analysis between malignant cells and the cancer-type-level normal reference using edgeR (57). The corresponding normal reference cell type for each cancer type is listed in Supplementary Table 11. Drug efficacy is then assessed using DrInsight (58), which identifies concordantly expressed genes against the L1000 dataset, computes connectivity via an outlier-sum approach, and evaluates significance with a Kolmogorov-Smirnov test (59) followed by Benjamini-Hochberg correction (60). Drug–subpopulation pairs with FDR < 0.05 are classified as drug responders, whereas those with FDR > 0.95 are classified as drug non-responders, enabling analysis of efficacy heterogeneity across malignant subpopulations. Drug–subpopulation pairs with 0.05 ≤ FDR ≤ 0.95 are treated as uncertain and excluded from responder–non-responder comparisons. The upper-tail cutoff follows previous studies using local FDR > 0.95 to define unchanged or unregulated features (61–63).

### 4.6. Integrated differential analysis of pharmacodynamic heterogeneity groups

To account for batch effects, we conduct differential analysis exclusively between effective and ineffective groups within the same project. Consequently, subsequent analyses focus on drugs associated with each cancer type that have at least one project containing both effective and ineffective groups. To aggregate multiple differential gene expression analysis results for a single drug, we utilize metaVolcanoR (v1.19.0) (64) to combine p-values and log-fold changes, denoting these as metaP and metaFC, respectively. For meta-analysis results, genes are ranked from highest to lowest based on the absolute product of log(metaP) and metaFC. When only a single differential gene analysis result is available, genes are ranked based on the absolute values of their log-fold changes in descending order.

### 4.7. Drug grouping based on pathway enrichment

To enable meaningful interpretation across more than 1,000 drugs, we categorize drugs based on their mechanisms of action. Using integrated differential gene expression results from the previous step, we apply Gene Set Enrichment Analysis (GSEA) (65) via ClusterProfiler (v4.16.6) (66) to identify significantly enriched Gene Ontology (GO) terms (67, 68) for each drug. Pathways with adjusted p-value < 0.05 are considered statistically significant. The NES for these pathways serve as feature vectors representing each drug. The enrichment results are presented in **Figure. S7**.

Subsequently, hierarchical clustering is performed on the NES matrix to group drugs according to their pathway-level similarities. We define meaningful drug clusters as those in which all constituent drugs share at least one consistently upregulated or downregulated pathway and which contain more than 5% of the total drug set. To determine the optimal cluster number, we search the clustering hyperparameter space across the range [2, N], where N represents the total number of drugs. The final cluster number is selected to maximize the total number of drugs assigned to meaningful clusters.

### 4.8. Identification of hub genes driving PharmITH

To identify key genes associated with each drug efficacy group, we integrate differential gene analysis results across drugs within the group. The votecount*_v_* method, implemented in the metaVolcanoR package, is applied to detect consistently differentially expressed genes using an absolute log2 fold-change threshold of 1 and default values for all other parameters. Subsequently, protein-protein interaction (PPI) networks are constructed for these consensus genes using the STRING database (v12.0) (69) with a high-confidence interaction score cutoff of 0.700. Top hub genes within each high-confidence PPI network are identified using the CytoHubba (v0.1) (70) plugin, which assesses node centrality based on 12 topological algorithms. Genes ranking among the top 20 in at least six of these algorithms are defined as hub genes.

### 4.9. Multi-omics-based heterogeneous gene regulatory network

To uncover transcription factors associated with cellular state heterogeneity and pharmacodynamic heterogeneity, we construct enhancer-driven gene regulatory networks (eGRNs) using SCENIC+ (16), which integrates single-cell ATAC-seq and RNA-seq data to link regulatory elements to target genes (Supplementary Note 1). We identify transcription factors connected to hub genes by analyzing differentially accessible regions (DARs) between phenotypically distinct clusters. To improve sensitivity for detecting transcription factor binding sites, the parameter dem_log2fc_thr is adjusted from the default value of 1.0 to 0.5. For samples with only scRNA-seq data available, we perform regulatory inference using pySCENIC (v0.12.1) (71), with results summarized in **Supplementary Table 8**.

### 4.10. Identification of heterogeneity-associated SNV events

We perform single-nucleotide variant (SNV) detection on scRNA-seq data using SComatic (72) with default parameters and the provided RNA editing file (73–75), beginning with conversion of raw FASTQ files into BAM format for downstream analyses. To explore SNVs potentially linked to heterogeneity, we focus on those uniquely detected within individual functional subpopulations. The identified SNVs are then annotated using ANNOVAR (76) and RefGene (77) for the annotation of genomic region. Fisher’s exact test is applied to identify mutation types significantly enriched in specific malignant subpopulations or drug response groups. To further investigate links between gene mutations and drug response, PPI networks are constructed connecting mutated genes to previously identified hub genes. This integrated approach enables investigation of PharmITH from the perspective of somatic mutations.

### 4.11. Common MPs identified via CCC in drug response heterogeneity

Cell-cell communication between distinct malignant cell subpopulations is analyzed using the R package LIANA+ (v1.4.0) (78, 79). The “Consensus” database is employed for ligand-receptor interaction analysis, with all processing steps using default parameters. A specificity rank threshold of 0.05 is set to ensure result significance. Analyses are conducted separately from the perspectives of malignant cell state heterogeneity and drug response heterogeneity (**Supplementary Table 9, Supplementary Table 10**).

### 4.12. Identification of heterogeneity-associated specific alternative splicing genes

To systematically identify PharmITH-associated alternative splicing events, we first align scRNA-seq raw reads (FASTQ) to the reference genome using STAR (v2.7.5a) (80) to generate BAM files. Compared to alternative approaches, the combined Splicing Z-score (SpliZ) (81) and SICILIAN (82) framework offers superior sensitivity in detecting splicing variations under low sequencing depth and avoids biases inherent in annotation-dependent tools. We therefore employ the SICILIAN pipeline on STAR-aligned outputs under default parameters to rigorously distinguish high-confidence splice junctions from false positives, providing a robust and annotation-free statistical framework. Leveraging SICILIAN’s junction count matrix, the SpliZ algorithm is subsequently applied along two axes—cell-state heterogeneity and drug-response heterogeneity—enabling identification of genes and splice variants specifically associated with each dimension and prioritization of high-confidence candidates for further investigation.

### 4.13. Knock-down with siRNA

Tumor cell lines A549 (LUAD) and MDA-MB-231 (TNBC) a are provided by Stem Cell Bank, Chinese Academy of Sciences (Shanghai, China). For knock-down (KD) of hub gene protein expressions, 0.8 μg of target siRNA or scrambled siRNA (Thermo Fisher Scientific, Silencer™ select non-targeting siRNAs) is mixed with 20 μL Lipo800 transfection reagent (Beyotime) in 1 mL transfection medium (without FBS) and incubates at room temperature for 30 min, and a mock control (no siRNA) is used as the Ctrl for the experiment. Following incubation, 300 μL of transfection medium containing the transfection complex is added to cells in 300 μL cell culture medium to achieve a final siRNA concentration of 0.4 μg/mL. After 24 h incubation at 37°C the transfection medium containing siRNA is replaced with fresh cell culture medium. Time 0 of the growth curve is defined as when the medium was replaced. The concentrations of inhibitors (ARP-100, 50 nM; PP2, 20 nM; Anticancer agent 195, 5 μM) were selected based on previous literatures and preliminary tests. CCK-8 kit is used to examine cell proliferation and viability, which was calculated based on the 450 nm absorbance, and Knock-down efficiency is validated by Western Blot analysis. Three independent replicates (n = 3) are set for each individual treatment or Ctrl group. One-way analysis of variance (ANOVA) and Turkey’s post-hoc tests were used to calculate the differences between groups. P values under 0.05 were considered statistically significant.

### 4.14. Western Blot of protein expression

Cell lysates are prepared using RIPA buffer and protein concentrations are determined by BCA assay. Samples are mixed with loading buffer, denatured at 100°C, and separated on polyacrylamide gels (20 μg protein/lane) by electrophoresis at 100 V in SDS-PAGE buffer on ice for approximately 2 h. Proteins are transferred onto PVDF membranes using a Bio-Rad semi-dry transfer system. Membranes are blocked with 5% skim milk in TBST overnight at 4°C and subsequently incubate with diluted primary antibodies for 1 h. HRP-conjugated secondary antibodies are then applied, followed by detection with BeyoECL Moon chemiluminescence reagent (Beyotime). Protein bands are visualized using a Tannon 5500 imaging system and band intensities are quantified with ImageJ v1.53. Three independent replicates (n = 3) are set for each individual group. One-way analysis of variance (ANOVA) and Turkey’s post-hoc tests were used to calculate the differences between groups. P values under 0.05 were considered statistically significant.

### 4.15. Mouse tumor model and inhibitor treatments

Male BALB/c nude mice (6 weeks old) are obtained from JSJ-Lab (Shanghai, China). All animal experiments are performed in accordance with protocols approved by the Institutional Animal Care and Use Committee (IACUC No. GZ2024-075). This study does not involve human participants, human samples, or identifiable human data. Following one week of acclimation in our animal facility, mice are subcutaneously injected with 2×10^6^ A549 cells or 1×10^6^ MDA-MB-231 cells in 100 μL cell culture medium mixed with Corning Matrigel matrix (3:1 ratio). When tumor volumes reached 100 mm^3^, mice are randomly allocated into different groups (n = 4) and administered daily intraperitoneal injections of various inhibitors (MedChemExpress) including ARP-100 (2 mg/kg/d), PP2 (1 mg/kg/d), TAS0612 (30 mg/kg/d), and Anticancer agent 195 (10 mg/kg/d) for two weeks. At the experimental endpoint, mice are euthanized by cervical dislocation and tumor tissues are harvested for subsequent analysis. One-way analysis of variance (ANOVA) and Turkey’s post-hoc tests were used to calculate the differences between groups. P values under 0.05 were considered statistically significant.

### 4.16. Academic writing refinement

The writing in portions of the article is polished using large language model.

## Data Availability

Main results and related functions are publicly available through a web interface at pharmith.net. To obtain the complete data, please contact the corresponding author.

## Acknowledgments

This work is supported by Research Funds of Joint Research Center for Occupational Medicine and Health of IHM [grant No. OMH-2024-041] and National Natural Science Foundation of China [grant No. 42177417]. This work is also funded by Natural Science Research Project for Excellent Young Scholars of Anhui Educational Committee [grant No. 2024AH030014] and Scientific Research Foundation for High-level Talents of Anhui University of Science and Technology [grant No. YJ20240007].

## Author contributions

Z.R. was involved in the conceptualization, supervision, methodology, and writing. Y.R. was involved in the conceptualization, methodology, and writing. Z.L. was involved in methodology, data curation. S.G. was involved in methodology. Y.X. was involved in data curation. H.X. was involved in conceptualization, supervision, funding acquisition and writing.

