## Supplementary Information for "Mechanistic Decoding of Pharmacodynamic Intratumoral Heterogeneity: A Pan-Cancer Regulatory Framework"

### List of contents

#### 1. Supplementary Notes


**Supplementary Note 1.** A pipeline for mapping enhancer-driven regulatory networks via integrated multi-omics

**Supplementary Note 2.** Enhancer-driven transcription factor networks shape malignant cell state heterogeneity

**Supplementary Note 3.** Transcription factor regulatory networks define drug group-specific response programs

**Supplementary Note 4.** Somatic variants associated with predicted drug-response states in the LUAD case study

**Supplementary Note 5.** Hub gene-centered intercellular communication heterogeneity within intratumoral populations

**Supplementary Note 6.** Ligand-receptor signaling defines an intercellular regulatory dimension of PharmITH

**Supplementary Note 7.** Alternative splicing defines a post-transcriptional regulatory dimension of PharmITH

**Supplementary Note 8.** Extended Pathway-level rewiring mechanisms underlying PharmITH


#### 2. Supplementary Figures


**Figure S1.** Molecular docking of ARP-100 with MMP2 and PP2 with FYN

**Figure S2.** Tumor growth and body weights of triple-negative breast cancer (TNBC) model

**Figure S3.** Mapping DARs of Regulatory Heterogeneity within Tumors

**Figure S4.** Knockdown of ELF3 in LUAD Cell Line

**Figure S5.** Original western blot images with molecular marker bands

**Figure S6.** Overview of cell types from scBERT

**Figure S7.** Overview of pathway enrichment across cancer drug groups

**Figure S8.** Deciphering the transcription factor-hub gene regulatory network in PharmITH subpopulations


#### 3. Supplementary Tables


**Supplementary Table 1.** Malignant meta-programs and corresponding functional gene sets

**Supplementary Table 2.** List of PharmITH hub genes

**Supplementary Table 3.** Case study of hub genes based on literature curation

**Supplementary Table 4.** Enhancer-informed regulatory network for hub genes in LUAD

**Supplementary Table 5.** Somatic mutations associated with PharmITH in LUAD

**Supplementary Table 6.** Context-specific communication networks of hub genes in LUAD

**Supplementary Table 7.** PharmITH-associated alternative splicing events in LUAD

**Supplementary Table 8.** Cancer-type-specific regulatory networks involving hub genes

**Supplementary Table 9.** Hub gene-mediated cell-cell communication in the tumor microenvironment

**Supplementary Table 10.** Differential cell-cell communication of drug-response malignant subpopulations

### Supplementary Notes

#### Note 1. A pipeline for mapping enhancer-driven regulatory networks via integrated multi-omics

To investigate the regulatory roles of transcription factors in malignant subpopulations and drug-response heterogeneity from a chromatin accessibility perspective, we employ SCENIC+ [1] to construct enhancer-driven gene regulatory networks (eGRNs). Specifically, we integrate single cell ATAC sequencing (scATAC-seq) and single cell RNA sequencing (scRNA-seq) datasets, align both modalities into a shared latent space, and annotate scATAC-seq cells using transcriptomic references. This approach generates a cell-resolved chromatin landscape that recapitulates the heterogeneous meta-programs (MPs) observed in the transcriptome.

The SCENIC+ workflow consists of three main steps: (1) identifying candidate enhancers by defining differentially accessible regions (DARs) from chromatin accessibility profiles; (2) detecting transcription factor binding motifs; (3) linking transcription factors to enhancers and their target genes to reconstruct eGRNs.

To align with our research objectives, we adapt the DAR analysis as follows:

(1) Intratumoral heterogeneity (ITH): We compute DARs across distinct malignant subpopulations to identify ITH-associated transcription factors, thereby capturing intra-tumoral differences in regulatory potential.

(2) Pharmacodynamic intratumoral heterogeneity (PharmITH): Within each sample, we classify malignant subpopulations as responsive or non-responsive clusters based on drug sensitivity. Responsiveness is determined using a representative score $S$ for each cluster within a given drug group, defined as:

$$S = E^2 - C^2$$

where $E$ denotes the number of effective drugs and $C$ the number of control drugs in the group. A positive score ($S > 0$) indicates sensitivity to the drug group, while a negative score ($S < 0$) indicates resistance or non-response. We rank clusters by $S$ within each drug group; the top three are designated as significantly responsive and the bottom three as significantly non-responsive. DARs between these groups are computed for subsequent transcription factor analysis. We then link the identified hub genes to these DARs to characterize response-associated transcription factor–enhancer–hub gene relationships.

**Note 2. Enhancer-driven Transcription factor networks shape malignant cell state heterogeneity**

Following the workflow in Figure. 3a, we use CisTopic to delineate MP-specific DARs in malignant cells and define representative enhancer segments. These regions provide a cis-regulatory coordinate system to relate transcription factor activity to hub genes. Based on these regions, SCENIC+ infers enhancer-driven triplets (transcription factor–enhancer–target), which we map to hub gene set.

Joint profiling of gene-based regulon specificity scores (RSS) and transcription factor expression across malignant MPs (Fig. 3b) identifies the top RSS-ranked transcription factors for each cellular state:

- **Epithelial.Senescence**: FOXA2 (+), FOXA1 (+), YBX1 (−)
- **Protein.maturation:** ATF5 (+), EHF (−), JUN (−), CREM (+)
- **Stress.LUAD.I:** YBX1 (−), MAFB (−), SPI1 (−)
- **Stress.LUAD.II:** ATF3 (−), CREM (−), MAFB (−)

Here, "+/−" indicates the direction of correlation between transcription factor expression and hub genes. Consistent with prior studies, FOXA1 and FOXA2 are lineage-defining and lineage-vulnerable factors in lung adenocarcinoma (LUAD) that support epithelial identity and differentiation [2]. ATF5 functions as a stress-survival regulator that enhances tumor cell persistence under proteostasis pressure via BCL-2 and MCL-1 [3–5], whereas ATF3 is a canonical stress-induced transcription factor implicated in metabolism, immunity, and tumorigenesis [6,7].

Ranking transcription factors by their frequency among the top 100 triplets reveals ELF3, YBX1, KLF5, SPI1, and MAFB as the most recurrent regulators, connecting 16, 10, 7, 6, and 5 hub genes, respectively (Fig. 3d). Their regulatory networks concentrate in distinct biological modules that together shape drug-response phenotypes:

**ELF3 — epithelial differentiation and adhesion coupled to protein maturation**
**Positive correlations:** CDH1, EPCAM, ICAM1, RPS3A
**Negative correlations:** AIF1, CD4, HCLS1, FCER1G, CD68, CXCR4, CD163, FN1, LCP2, ITGB2, CD44, APOE, IL1B
The ELF3 network links epithelial adhesion and identity genes (CDH1, EPCAM, ICAM1) to immune-metabolic and myeloid markers (APOE, FCER1G, CD68, CD163), suggesting coupling between epithelial state and protein maturation or proteostasis programs. Notably, in EGFR-TKI-resistant settings, ELF3 inhibition induces tumor cell death and improves responses to EGFR-targeted therapy [8]. More recently, ELF3 has been shown to restrain ferroptosis and thereby promote lung tumorigenesis, providing a mechanistic link to stress survival and drug tolerance [9].

**YBX1 — autophagy-linked stress survival and reduced platinum sensitivity**
**Positive correlations:** TYROBP, CD68, FCER1G, HCLS1, IL1B, APOE, CSF1R
**Negative correlations:** CCL5, EPCAM, IL2RB
YBX1 correlates positively with myeloid and inflammatory nodes (TYROBP, CSF1R, IL1B, CD68) and negatively with the epithelial marker EPCAM and the T-cell signaling gene IL2RB, consistent with a stress-adapted, less epithelial state. Functionally, YBX1 enhances autophagy via the p110β–Vps34–Beclin1 axis and reduces cisplatin sensitivity, aligning with the associations observed here [10].

**KLF5 — epithelial plasticity, EMT, and DNA-damage response**
**Positive correlations:** ICAM1, FN1
**Negative correlations:** TYROBP, AIF1, HCLS1, CD68, FCER1G
Enrichment for matrix–adhesion and migration genes (FN1, ICAM1) indicates EMT-related plasticity. Experimentally, KLF5 suppression markedly sensitizes NSCLC cells to cisplatin-induced apoptosis and attenuates the γH2AX–Chk1/Chk2 phosphorylation cascade, thereby limiting DNA repair and overcoming chemoresistance—consistent with the "EMT–plasticity ↔ drug response" axis in our network [11].

**SPI1 (PU.1) and MAFB — myeloid and tumor-associated macrophage programs**
**SPI1 positive correlations:** TYROBP, CD68, FN1, CD163, CD4
**MAFB positive correlations:** CD163, CXCL8, FCER1G, IL1B
**Shared negative correlation:** CD3E
Both transcription factors align with tumor-associated macrophage (TAM) signatures (CD68, CD163) and chemokine axes (CXCL8), indicating strong coupling to the myeloid microenvironment. Mechanistically, SPI1 activates CCL7, promoting M2 macrophage accumulation and dissemination [12], whereas MAFB fosters M2-like polarization [13]. In lung cancer, TAMs are linked to tumor growth, angiogenesis, metastasis, and resistance to therapy. Combinatorial TAM-targeting with chemotherapy or immunotherapy has shown clinical benefit, consistent with the myeloid intervention points highlighted by our network [14].

The enhancer-driven regulatory network in Fig. 3d converges on three cooperating axes: epithelial differentiation and protein maturation (ELF3, KLF5), stress survival and drug tolerance (YBX1), and myeloid inflammatory microenvironment (SPI1, MAFB). Together, these axes map onto hub genes and support a model in which tumor-intrinsic programs and microenvironmental remodeling jointly shape distinct drug-response states. Notably, the state-specific nature of enhancer accessibility implies that the same hub gene can be under different transcription factor control in different cellular states, providing a regulatory basis for the observed heterogeneity in drug responses across malignant MPs.

**Note 3. Transcription factor regulatory networks define drug group-specific response programs**

**Drug group DG1**

DG1 comprises 99 approved or investigational agents spanning diverse classes, including EGFR/ERBB inhibitors (erlotinib, lapatinib), BCR–ABL/Src inhibitors (dasatinib, bosutinib), a BRAF inhibitor (vemurafenib), endocrine modulators (tamoxifen, spironolactone, progesterone), glucocorticoids (betamethasone, methylprednisolone, clobetasol), antimetabolites and alkylators (fluorouracil, floxuridine, temozolomide), antivirals (entecavir, indinavir, raltegravir, nelfinavir), anti-infectives (amikacin, cefixime, ceforanide, nafcillin), PPARγ agonists (pioglitazone, rosiglitazone), and DPP-4 inhibitors (linagliptin, sitagliptin).

Drug–response profiling indicates pronounced sensitivity in Protein.maturation and Stress.LUAD.I states, whereas Epithelial.Senescence in control samples displays minimal or inverse activity. ATAC-seq identifies 34,011 accessible regions in responders versus 5,713 in control samples, with fewer than 280 shared peaks (≥1-bp overlap), underscoring state-specific chromatin remodeling.

Using SCENIC+, we identify responder-preferred transcription factors — SPDEF (−), EHF (−), SPI1 (+), ELF3 (−), and KLF5 (−) — based on responder-biased RSS deltas. Integrating transcription factor→enhancer connections (eRegulon_100) with enhancer→gene links yields enhancer-centered subnetworks in which EHF (12 genes), SPI1 (11 genes), SPDEF (7 genes), ELF3 (7 genes), and KLF5 (6 genes) emerge as the most recurrent regulators of DG1 efficacy genes (Fig. 3d).

These transcription factors bifurcate into two functional arms: an immune–adhesion arm (SPI1-linked CSF1R, IL1B, ITGB2, ITGAX, TYROBP, CXCR4) [15] and an epithelial arm (EHF/ELF3/SPDEF/KLF5-linked CDH1, and EHF/SPDEF-linked EGFR) [16–18], with the former showing broader gene coverage. Consistent with this regulatory architecture, DG1 contains direct modulators of these modules, including erlotinib and lapatinib (targeting EGFR/ERBB2) [19,20] and plerixafor (targeting CXCR4) [21].

Notably, dasatinib and bosutinib inhibit Src-family kinases (SFKs) that act immediately downstream of integrin engagement with the extracellular matrix (ECM) through the FAK/Src axis, thereby dampening focal-adhesion signaling, cytoskeletal remodeling, and cell migration—a mechanism consistent with the ITGB2/ITGAX–ECM adhesion program captured in the immune–adhesion arm [22–24]. In parallel, PPARγ agonists (pioglitazone and rosiglitazone) promote anti-inflammatory remodeling of myeloid and chemokine modules, including IL1B and CCL2 contexts [25,26]. Taken together, DG1 response patterns are organized around immune-receptor and adhesion programs with embedded epithelial nodes, linking the inferred transcription factor circuitry to the established pharmacology of specific drug classes.

**Drug group DG4**

DG4 comprises mechanistically diverse agents, including ErbB/ALK pathway inhibitors (afatinib, alectinib, crizotinib) [27,28], DNA–topoisomerase poisons (etoposide, teniposide, topotecan) [29,30], microtubule disruptors (vincristine, vinblastine, vinorelbine, paclitaxel) [31], an epigenetic modulator (romidepsin, a class I HDAC inhibitor) [32], and a Hippo–YAP pathway blocker (verteporfin) [33]—consistent with established targets for these compounds.

Drug–response profiling reveals pronounced sensitivity in Epithelial.Senescence subpopulation, whereas control MPs (Stress.LUAD.I/II, Protein.maturation) show weak or opposite activity. The DG4 efficacy gene set is enriched for a coherent immune–adhesion axis, including TNF, IL1B, CXCL8, ITGB2, CXCR4, CD44, PTPRC/CD45, FCGR3A, TYROBP, CD86, and CCL5, together with macrophage and monocyte markers (CD68, CD163), T-cell activation nodes (CD2, CD4, LCK), and an embedded epithelial node (CDH1, with APOE).

Enhancer-resolved eRegulon mapping highlights recurrent regulators that bridge epithelial and myeloid circuitry—ELF3, SPDEF, KLF5, ETS2, and SPI1/PU.1—consistent with SCENIC+ inference of enhancer-driven gene regulatory networks (GRNs) and prior functional assignments of these transcription factors to epithelial identity (ELF3, KLF5, SPDEF) and myeloid programming (SPI1), with ETS2 linking inflammatory responses [34].

The greater negative connectivity in responsive MPs is consistent with the directionality of DG4 pharmacodynamics. A subset of DG4 agents—including immunogenic cell death (ICD)-inducing anthracyclines [35], TOP1/TOP2 poisons that trigger cGAS–STING–NF-κB/IRF signaling [36,37], and the HDAC inhibitor romidepsin—converge to upregulate chemokine, adhesion, and interferon-response modules, thereby countering the immune-low, epithelial-high baseline of Epithelial.Senescence more effectively than in stress-maturation control states.

**Drug group DG5**

DG5 comprises mechanistically heterogeneous agents with a prominent immunomodulatory core, including multiple glucocorticoids (dexamethasone, budesonide, betamethasone, prednisolone, halcinonide, amcinonide, loteprednol, medrysone), mTOR pathway inhibitors (sirolimus, temsirolimus), MEK inhibitors (trametinib, selumetinib), an EGFR inhibitor (gefitinib), multikinase TKIs (sorafenib, axitinib), a BTK inhibitor (ibrutinib), a calcineurin inhibitor (tacrolimus), a CDK4/6 inhibitor (palbociclib), PDE5 inhibitors (sildenafil, tadalafil), and endocrine modulators (tamoxifen, toremifene, bicalutamide, finasteride).

Drug–response profiling indicates greater activity in Stress.LUAD.II and Protein.maturation MPs, whereas Epithelial.Senescence and Protein.maturation programs in the control set show weaker or opposite responses.

The DG5 efficacy gene set consolidates an immune–adhesion and T-cell activation axis, including PTPRC, FCGR3A, ITGB2, IL1B, CCL5, FCER1G, TYROBP, CD2, CD3D, CD3E, LCP2, HCLS1, AIF1, CD68, and CXCR4, with embedded epithelial nodes (EPCAM, CDH1). The enhancer-resolved eRegulon for DG5 forms a compact, immune-adhesion-centric network in which ELF1 functions as the principal hub, with SPI1, CREM, and ETS2 scaffolding myeloid and T-cell receptor (TCR)/inflammatory signaling, whereas ELF3 provides only weak coupling to epithelial programs. Network connections converge on T-cell and myeloid activation, receptor transduction through the Fc-receptor–DAP12 axis, integrin adhesion, and chemotaxis, with limited epithelial embedding occurring primarily via EPCAM.

Mechanistically, CDH1 and EPCAM exhibit higher RSS in control cells, whereas LCP2, IL1B, TYROBP, ITGB2, HCLS1, RPS27A, RPS3A, CXCR4, FCER1G, CD68, AIF1, and CD2 show higher RSS in responsive cells. Consistent with these state differences, glucocorticoids broadly suppress pro-inflammatory mediators [38] and dampen TLR-driven signaling, tacrolimus blocks calcineurin–NFAT-dependent T-cell activation [39], and sirolimus and temsirolimus modulate T-cell programs through mTOR inhibition [40], whereas MEK inhibition remodels myeloid and chemokine outputs in the tumor microenvironment. Together, these pharmacological mechanisms are consistent with the stronger reversal of the immune-high signature observed in responsive MPs compared to Epithelial.Senescence-skewed control states.

**Note 4. Somatic variants associated with predicted drug-response states in the LUAD case study**

To examine somatic variation as an additional regulatory-context layer, we applied SComatic [41] to infer candidate variant sites from scRNA-seq raw reads across five LUAD samples, identifying 8,549 variant events. Fisher's exact tests identified 41 events associated with predicted responder subpopulations across three drug groups, including 13 events in DG1, 4 in DG2, and 24 in DG3. None of the corresponding genes overlapped with the predefined PharmITH-associated hub genes. We therefore examined whether selected variant backgrounds were accompanied by differences in hub-gene activity or pathway states. These analyses provide associative evidence and do not establish that the identified variants causally determine drug response. Two representative examples involving FAT1 and TCIRG1 are described below, and the complete results are provided in Supplementary Table 5.

FAT1-variant cells showed increased activity of YAP/TEAD transcriptional and Hippo-pathway-related modules, including CTGF, CYR61, ANKRD1, AMOTL2, BIRC5, and AURKA. This expression pattern is consistent with previous evidence that FAT1 loss of function can be associated with YAP/TAZ activation [42]. In the corresponding drug group, the FAT1-variant state was also associated with a higher predicted response propensity to the YAP–TEAD inhibitor verteporfin [43,44]. These observations suggest a candidate association among the FAT1 variant background, YAP/TEAD pathway activity, and predicted verteporfin response.

TCIRG1-variant cells showed increased activity of lysosomal-acidification, TFEB-regulated, and autophagy-related gene modules. Because TCIRG1 encodes a V-ATPase subunit involved in lysosomal acidification, these expression patterns indicate an association between the TCIRG1 variant background and a lysosome–autophagy-related cellular state. This state was accompanied by changes in EGFR/ERBB2 signaling, adhesion-related genes including FN1 and ITGB2, and inflammatory mediators including IL1B, CXCL8, and CCL2. The corresponding cells also showed higher predicted response propensities to lysosome- or autophagy-modulating compounds such as chloroquine and tamoxifen [45,46].

**Note 5. Hub gene-centered intercellular communication heterogeneity within intratumoral populations**

As shown in Figure. 4a, cell-cell communication analysis reveals distinct signaling niches among the four malignant LUAD subpopulations. The Epithelial.Senescence cluster shows a pronounced senescence-associated secretory phenotype (SASP), evidenced by enriched JAK–STAT cytokine signaling and frequent interactions with cytotoxic lymphocytes ($CD8^+$ T cells and NK cells) and myeloid cells. These senescent-like tumor cells likely secrete inflammatory mediators (e.g., IL-6, IL-8) that activate STAT3 and attract immune effectors, consistent with SASP-driven immunosurveillance attempts. Notably, EGFR-pathway signaling remains active in this cluster, suggesting that even growth-arrested malignant cells retain growth factor receptor engagement.

In contrast, the Protein.maturation subpopulation engages predominantly with monocytes and fibroblasts, aligning with its higher utilization of Wnt and TGF-β pathways. This cluster shows strong CD44-mediated interactions with extracellular matrix (ECM) components: for instance, perivascular stromal cells (pericytes) provide collagen ligands (COL1/4) that bind CD44 on Protein.maturation cells. Such contacts suggest a more mesenchymal, matrix-attached phenotype embedded in a stromal and pro-survival signaling context. Two stress-adapted LUAD subpopulations (I and II) exhibit robust autocrine and paracrine signaling in the tumor microenvironment by secreting high levels of CXCL8 (IL-8) and CCL5 that engage chemokine receptor ACKR1 on perivascular endothelial cells. This concentrates chemokines perivascularly, shaping immune cell trafficking: IL-8 recruits neutrophils and MDSCs, dampening cytotoxic immunity and promoting EMT.

Stress.LUAD cells also upregulate inflammatory pathways (NF-κB, TNFα), indicating an immunosuppressive, inflammation-driven phenotype. Both I and II express integrin β2 (ITGB2) that binds THY1 on pericytes. Stromal cells provide ECM signals (collagens to CD44 and MYL9 to CD69 on II), defining a distinct stromal-interaction context around these malignant states. NK cells secrete SPON2 that binds ITGB2 on II, indicating tumor-innate immune crosstalk. Pathway biases differ: II is enriched for NF-κB/TNFα-related signaling, whereas I shows EGFR/JAK-STAT/TGF-β-related signaling. Notably, TGF-β enrichment in I suggests an EMT-promoting, immunosuppressive microenvironment. Consistently, we observe upregulated NF-κB and TNF inflammatory signaling in Stress.LUAD cells, along with significant crosstalk with immunosuppressive myeloid cells, such as alveolar macrophages. Collectively, these findings demonstrate that LR pairs involving hub genes define distinct communication landscapes across malignant subpopulations, linking PharmITH-associated hub genes to distinct intercellular signaling contexts across malignant states.

**Note 6. Ligand-receptor signaling defines an intercellular regulatory dimension of PharmITH**

Investigating cell-cell crosstalk through the lens of treatment response identifies substantial overlap between ITH-associated and PharmITH-associated communication patterns. Five ligand-receptor pairs involving hub genes show divergent signaling profiles between predicted responder and non-responder subpopulations. Together, these interactions define an intercellular regulatory dimension of PharmITH.

**Distinct LR interactions in drug-responder malignant subpopulations**

**FN1-CD79A**: In drug-responder tumors, the FN1-CD79A LR pair is uniquely identified. Malignant cells upregulate FN1 and interact directly with CD79A$^{+}$ B cells. Although elevated fibronectin is commonly linked to chemoresistance through integrin binding and activation of pro-survival pathways such as YAP and SOX2 [47], its specific interaction with CD79A in responder tumors suggests an alternative, immune-favorable role. Fibronectin can support immune niches; for example, fibronectin-integrin signaling inhibits apoptosis in B-cell malignancies [48]. In solid tumors, an FN1-rich microenvironment may promote the formation of tertiary lymphoid structures or enhance B-cell retention. Notably, the presence of CD79A$^{+}$ B cells and tertiary lymphoid structures correlates strongly with improved therapy response. Thus, the preferential FN1-CD79A interaction in responder subpopulations is consistent with an immune-supportive ECM context and links this hub-gene interaction to the responder-associated communication landscape [47,48].

**Distinct LR interactions in drug non-responder malignant subpopulations**

By contrast, two LR pairs unique to the drug non-responder subpopulation, ICAM1-MUC1 and TFF3-CXCR4, exemplify how ITH can foster a pro-survival, drug non-responder phenotype. These interactions involve hub genes associated with aggressiveness.

**ICAM1-MUC1:** In non-responder tumors, subpopulations expressing elevated levels of ICAM1 engage with MUC1 present on adjacent cells. This interaction enhances malignant phenotypes; ICAM1-MUC1 binding reprograms the tumor microenvironment through altered secretion of cytokines and proteases and further upregulates ICAM1 expression in cancer cells, ultimately promoting cell migration and metastasis [49]. MUC1 is a well-established oncoprotein that drives therapy resistance and is frequently overexpressed in epithelial cancers. It confers multi-drug resistance through several mechanisms including reduced uptake of chemotherapeutic agents such as 5-FU, activation of survival pathways such as STAT3 and HIF1α, and metabolic reprogramming that together enhance chemo- and radio-resistance [50]. Thus, the preferential ICAM1-MUC1 interaction places non-responder subpopulations within an adhesion- and survival-associated communication context consistent with known MUC1-linked resistance biology.

**TFF3-CXCR4:** In non-responder tumors, Trefoil Factor 3 is secreted by cell types such as lymphatic endothelial cells and binds to CXCR4 present on cancer cells. This LR interaction promotes pro-tumorigenic behaviors. TFF3 is highly upregulated in advanced cancers and has been directly linked to therapy resistance and metastasis. For example, elevated TFF3 mediates resistance to endocrine and HER2-targeted therapies in breast carcinoma by enhancing cancer cell survival, stem-like properties, and invasiveness [51]. Mechanistically, TFF3 signals through CXCR4 and its coreceptor CXCR7 to stimulate cell migration and invasion [52]. In vitro studies confirm that TFF3 binding to CXCR4 and CXCR7 enhances cell motility via mechanisms independent of canonical ERK signaling, indicating a unique chemokine-like activity. Additionally, recent studies show that TFF3 also binds the receptor CD147 and activates downstream SRC/STAT3 and COX-2 signaling, resulting in increased prostaglandin E2 production that fuels colorectal cancer progression and metastasis [52]. Within the non-responder subpopulation, preferential TFF3-CXCR4 signaling is consistent with a motility-, stemness-, and survival-associated communication context supported by the established biology of TFF3/CXCR4 signaling [53]. Therefore, the preferential ICAM1-MUC1 and TFF3-CXCR4 interactions define an adhesion-, survival-, and motility-associated communication program enriched in predicted non-responder subpopulations.

**Differences in shared LR interactions between responder and non-responder malignant subpopulations.**

**CXCL8-ACKR1:** The chemokine interaction CXCL8-ACKR1 shows a marked response-associated difference in inflammatory communication. In drug group 1, a marked disparity in CXCL8-ACKR1 signaling is observed between drug-responder and drug non-responder subpopulations. CXCL8 is a pro-inflammatory and pro-angiogenic chemokine frequently upregulated in therapy non-responder tumors. ACKR1 acts as a decoy receptor that binds and sequesters CXCL8 without initiating signaling. In responder tumors, higher ACKR1 expression on endothelial cells helps neutralize excess CXCL8, thereby limiting its bioavailability. This scavenging function is generally associated with improved clinical outcomes and reduced metastasis. By contrast, non-responder tumors exhibit stronger CXCL8-ACKR1 engagement, suggesting either overwhelming CXCL8 production or loss of functional ACKR1 suppression. Consequently, the non-responder subpopulation experiences sustained CXCL8 signaling, which promotes cancer cell proliferation, stemness, and chemoresistance. Elevated CXCL8 recruits CXCR2-positive myeloid cells and activates survival pathways such as STAT3, fostering an immunosuppressive microenvironment that diminishes drug efficacy. Thus, the differential activity of the CXCL8-ACKR1 axis between responder and non-responder groups highlights an inflammatory signaling axis associated with predicted response heterogeneity.

**CXCL8-CD79A:** The CXCL8-CD79A interaction remains consistent between drug-responder and drug non-responder groups in multiple drug groups, indicating it does not contribute to differential treatment efficacy. This pair reflects IL-8 signaling in the context of B cells, marked by CD79A. Similar levels of IL-8 and B-cell presence across both subgroups suggest that neither B-cell infiltration nor IL-8 production were skewed by ITH in a way that influences drug response. Biologically, IL-8 primarily signals through CXCR1/2 receptors on neutrophils and myeloid cells rather than directly engaging B cells. Thus, the unchanged CXCL8-CD79A axis likely represents a baseline inflammatory milieu common to both phenotypes. Its comparable activity between groups suggests that this interaction represents a shared inflammatory communication feature rather than a major discriminator of predicted response states.

### Note 7. Alternative splicing defines a post-transcriptional regulatory dimension of PharmITH

Alternative splicing analysis reveals eight hub genes—DDN-AS1, FAM228B, CXADR, DIPK1A, LIX1L-AS1, PABPC1, STT3A-AS1, and PCED1B—with divergent splicing patterns between predicted drug-responder and drug non-responder subpopulations, revealing a post-transcriptional regulatory dimension of PharmITH.

#### Drug responder-associated splicing features

DDN-AS1 is a long non-coding antisense RNA at the DDN/PRKAG1 locus. While its role in LUAD remains unclear, it has been noted as upregulated in other cancers where it acted as an oncogenic driver [54]. In the LUAD context, DDN-AS1 splicing and expression were higher in drug-responder tumors than non-responder ones, although no direct studies connect this lncRNA to drug response. Loss of DDN-AS1 in non-responder cells might alter regulatory networks such as microRNA sponging or epigenetic modulation that confer a survival advantage, but further research is needed to confirm any role in therapy sensitivity.

FAM228B encodes a protein of largely unknown function. There is currently no known research linking FAM228B alternative splicing to drug response in LUAD. Its higher splicing inclusion in responder cases suggests it might be downregulated or functionally lost in non-responder tumors. Given the lack of functional studies, it remains speculative whether FAM228B contributes to drug sensitivity; it may simply be a passenger biomarker. Further investigation is required to determine if FAM228B isoforms influence cell survival or drug metabolism.

CXADR, also known as the coxsackievirus and adenovirus receptor, is a cell adhesion receptor often downregulated in more aggressive cancers. Higher CAR expression has been associated with greater responsiveness to certain treatments. Overexpression of CXADR was shown to increase glioma cell sensitivity to the chemotherapeutic paclitaxel [55]. In lung cancer cells, elevated CAR facilitates adenovirus-based therapies: non-responder A549 cells with higher CAR were more efficiently infected and killed by oncolytic adenovirus combined with cisplatin, illustrating that high CXADR levels can improve treatment efficacy [56]. The finding that CXADR splicing levels are higher in drug-responder LUAD cases aligns with these observations. Tumors retaining higher CAR expression and isoforms may be less malignant and more susceptible to therapy, whereas non-responder tumors often show CAR loss or aberrant splicing that reduces its function.

**Drug non-responder-associated splicing features**

DIPK1A, also known as FAM69A, is a transmembrane protein localized to the endoplasmic reticulum that undergoes alternative splicing into multiple isoforms. There is no direct literature linking DIPK1A to chemoresistance in LUAD; however, its increased splicing and expression in non-responder tumors suggests it could play a role in stress adaptation. Being an ER-localized protein, DIPK1A might influence protein folding or secretory pathways. Upregulation of DIPK1A isoforms in drug non-responder cells could hypothetically enhance the unfolded protein response or other survival pathways, thereby conferring tolerance to drugs. This remains conjectural in the absence of specific studies, but the differential splicing indicates DIPK1A as a candidate for further investigation in resistance mechanisms.

STT3A-AS1 is an antisense lncRNA to STT3A, a gene encoding the catalytic subunit of the oligosaccharyltransferase complex critical for N-linked glycosylation. While STT3A-AS1 itself has not been well characterized, its elevated splicing in non-responder LUAD cases likely reflects changes in the STT3A pathway. STT3A is frequently overexpressed in LUAD and is associated with worse prognosis [57]. STT3A activity promotes tumor cell survival and proliferation by enhancing glycosylation of key proteins and activating growth signals through MAPK and PI3K/AKT pathways. Knocking out or inhibiting STT3A can suppress LUAD cell growth and motility [57]. Higher STT3A-AS1 in non-responder tumors could indicate upregulation of STT3A and its pro-tumor effects. The lncRNA may stabilize STT3A mRNA or chromatin, or serve as a surrogate marker of STT3A expression. The net result would be a more aggressive, drug-tolerant phenotype. Although direct evidence tying STT3A-AS1 to drug response is lacking, the link between STT3A overexpression and therapy resistance is plausible, given that aggressive, STT3A-high tumors could better withstand anticancer treatments.

LIX1L-AS1 is the antisense transcript of LIX1L. Increased splicing of LIX1L-AS1 in non-responder LUAD parallels known roles of the LIX1L protein in cancer. Recent studies show LIX1L acts as a pro-metastatic, mesenchymal marker and contributes to therapy resistance. LIX1L is markedly upregulated during epithelial–mesenchymal transition in NSCLC and enhances cell migration, anoikis resistance, and notably confers resistance to EGFR tyrosine kinase inhibitors [58]. Mechanistically, LIX1L was found to bind nucleolin and promote ribosomal RNA synthesis, driving proliferation and EMT. Targeting this pathway can reverse EGFR-TKI resistance [58]. The antisense LIX1L-AS1 might be co-expressed with LIX1L or regulate its expression. Higher LIX1L-AS1 in drug non-responder LUAD likely indicates activation of the LIX1L axis, aligning with the known ability of LIX1L to promote drug non-responder, mesenchymal phenotypes. This makes the LIX1L/LIX1L-AS1 locus a potential contributor to TKI or chemotherapy resistance in LUAD, although the lncRNA's exact function remains to be elucidated.

PCED1B, also known as FAM113B, is a hydrolase enzyme that modifies cell-surface biopolymers and is subject to alternative splicing into multiple isoforms. Its role in lung cancer drug response has not been characterized in the literature. The observation that PCED1B splicing is higher in drug-responder LUAD and reduced in non-responder cases could imply that loss or altered splicing of PCED1B accompanies the development of resistance. Since PCED1B is associated with cell surface modifications, certain PCED1B isoforms might facilitate drug uptake or immune recognition, and their reduction in non-responder tumors helps cells evade these effects. However, no studies to date directly link PCED1B to chemosensitivity. It remains an interesting candidate: differences in PCED1B splicing between responder and non-responder tumors warrant further research to determine if this gene influences drug responsiveness or is merely a correlative biomarker.

PABPC1 is a pivotal RNA-binding protein involved in mRNA stabilization and translation, often upregulated in rapidly proliferating cancer cells. Paradoxically, drug-responder LUAD had higher PABPC1 splicing expression than non-responder tumors. Generally, PABPC1 overexpression is associated with aggressive tumor behavior and even therapy resistance. In HER2-positive breast cancer cells, the lncRNA SNHG14 drives up PABPC1 expression via epigenetic activation, leading to Nrf2 pathway activation and resistance to trastuzumab. Studies have proposed that aberrant PABPC1 activity can promote survival signals and DNA repair, contributing to multi-drug resistance [59]. The higher PABPC1 in responder LUAD patients might indicate that those tumors were more proliferative and hence initially responsive to cytotoxic drugs, whereas the non-responder tumors downregulated PABPC1 as an adaptation to reduce growth and evade drug action. Different PABPC1 splice isoforms might play distinct roles; non-responder cells might favor an isoform with reduced translational activity. While the direct causal link remains unproven in LUAD, the correlation underscores PABPC1's importance in cancer cell viability and suggests that changes in its splicing and expression can influence how tumors respond to treatment.

In summary, differential splicing patterns in the above genes correlate with drug sensitivity differences in LUAD. Genes with higher expression or inclusion in drug-responder tumors tend to be those whose loss or aberrant splicing is associated with a more aggressive, non-responder phenotype. Conversely, genes elevated in drug non-responder tumors are linked to pathways that promote survival under therapeutic stress. These findings are consistent with the notion that alternative splicing contributes to drug resistance in LUAD. However, for many of these specific genes, direct functional evidence is still lacking. Further research is needed to determine whether these splicing changes actively drive resistance or serve as biomarkers of other resistance mechanisms. Understanding these splicing–drug response relationships could inform new strategies, such as splicing-modulating therapies or biomarker-driven treatment selection, to overcome drug resistance in LUAD.

**Note 8. Extended pathway-level convergence of multi-layer regulatory signals in PharmITH**

To further characterize the pathway-level organization of the PharmITH regulatory landscape, we integrate regulatory features identified across transcription factor regulation, somatic variation, cell-cell communication, and alternative splicing according to their canonical pathway contexts. This pathway-centered analysis highlights recurrent functional modules shared across different classes of regulatory evidence and provides an expanded view of the six core pathways summarized in Figure. 5.

**Predicted drug responder-associated pathways**

**Wnt signaling pathway**

The epithelial cell adhesion molecule EPCAM functions as a hub gene and serves as a pivotal link between cell-cell adhesion and Wnt signaling in LUAD. Regulatory network data indicate that EPCAM expression is downregulated in the drug-responder malignant subpopulation, concomitant with upregulation of mesenchymal and immune genes. EPCAM is known to potentiate canonical Wnt/β-catenin signaling, as the cleaved EPCAM intracellular domain co-translocates with β-catenin to the nucleus and upregulates Wnt target genes [60]. Active Wnt signaling sustains cancer stemness and therapy resistance. These observations place the response-associated variation in EPCAM within an epithelial-plasticity and Wnt-related regulatory context. Alternative splicing events in Wnt pathway components, such as the switch between CD44 standard and variant isoforms, are also known to regulate epithelial-mesenchymal transition and impact drug responses [61].

Consistent with the main-text network, additional transcriptional, communication, and genetic features identified in responder states map to the same epithelial adhesion/Wnt axis. ELF3-CDH1 regulation provides a transcriptional component, LRP5- and PTPRM-associated communication provides a cell-surface signaling component, and FAT1/PTPRF variation adds genetic context. Together, these features identify Wnt/β-catenin signaling as a recurrent multi-layer regulatory module associated with predicted responder states rather than as the consequence of a single regulatory event.

**PI3K/AKT/mTOR signaling pathway**

In drug-responder LUAD subpopulations, the hub gene ICAM1 is embedded within a PI3K/AKT/mTOR-associated regulatory network involving growth-factor and survival signaling. ICAM1-fibrinogen ligation preserves tyrosine phosphorylation and recruits SHP-2, driving PI3K/AKT activation [62] that converts PIP2 to PIP3 and enables AKT phosphorylation at T308

by PDK1 and S473 by mTORC2 [63,64], ultimately conferring survival signaling in malignant cells.
The somatic-variation layer provides additional features associated with this pathway. HSP90AA1 variation occurs in a molecular context in which HSP90 is known to stabilize multiple pathway components, including AKT, mTOR, and EGFR, while HSP90 inhibition can suppress PI3K/AKT/mTOR signaling by destabilizing these proteins [65]. IRF2BPL variation provides an additional transcriptional-stress feature intersecting with PI3K/AKT-related programs.
Alternative splicing provides a further regulatory dimension. ICAM1 maintains constitutive expression, while SOD2 exhibits splicing heterogeneity within an oxidative-stress context linked to AKT/FOXO and mTOR-regulated cellular metabolism [64]. PRIMA1 demonstrates both mutation and alternative splicing, adding another genetic and post-transcriptional feature to this pathway-associated network.
Collectively, ICAM1-associated signaling, HSP90AA1 and IRF2BPL variation, and response-associated splicing features converge on a PI3K/AKT/mTOR-centered module in predicted responder subpopulations. The representation of this pathway across several regulatory layers supports its prioritization as a core component of the responder-associated PharmITH landscape.

**Predicted drug non-responder-associated pathways**

**NF-κB inflammatory pathway**

Tumor necrosis factor (TNF) is a central hub in a drug non-responder LUAD subpopulation, placing it within a prominent pro-inflammatory NF-κB-associated regulatory network. Its expression is associated with transcriptional regulators such as ATF5 and AP-1. Autocrine TNF-α production can persistently activate the canonical NF-κB pathway and reinforce tumor-cell survival [66], accompanied by expression of inflammatory chemokines including CXCL8 and CCL5.
The communication layer further highlights TNF- and CXCL8-associated inflammatory interactions involving malignant and microenvironmental populations. Previous studies in lung cancer have shown that neutrophil-associated inflammatory signaling can alter EGFR-TKI response [67], while TNF-α/NF-κB signaling has been directly implicated in acquired resistance in EGFR-mutated lung cancer [68]. NF-κB activation can also alter RNA-splicing patterns of responsive genes, providing a link to post-transcriptional regulation. At the same time, acute TNF-α exposure can enhance chemotherapy-induced cell death in specific contexts [69], underscoring the context-dependent nature of this signaling axis.
In our analysis, the recurrence of TNF/NF-κB-related transcriptional, intercellular, and post-transcriptional features defines an inflammatory module enriched in predicted non-responder states. Its agreement with established NF-κB-associated resistance mechanisms in NSCLC provides additional biological support for this pathway-level convergence.

**Lysosomal autophagy pathway**

A distinct drug non-responder LUAD subpopulation is characterized by high expression of the lysosomal protein LAPTM5, which functions as a hub gene. This pattern identifies autophagy-lysosomal activity as a prominent component of the predicted non-responder regulatory landscape. Consistently, gene regulatory data show that transcription factors such as ATF5 and MAFB are associated with lysosomal and myeloid genes including CD68 and CD163.

The somatic-variation layer independently supports the same pathway. TCIRG1-variant cells show increased activity of lysosomal-acidification, TFEB-regulated, and autophagy-related modules and display altered predicted responses to lysosome- or autophagy-modulating compounds including chloroquine and tamoxifen [45,46]. Thus, TCIRG1 variation provides a genetic feature that converges with the transcriptional and hub-gene evidence on lysosomal-autophagy regulation.

Functionally, LAPTM5 fortifies the lysosomal membrane and sustains autophagic flux, thereby enabling cells to survive chemotherapeutic stress. In NSCLC, LAPTM5 confers cisplatin resistance by preventing lysosomal membrane permeabilization and cell death [70]. Specifically, LAPTM5 inhibits the ubiquitination and degradation of LAMP1, preserving lysosomal integrity and blocking cathepsin release that would otherwise trigger apoptosis.

Together, LAPTM5-centered hub-gene activity, ATF5/MAFB-associated transcriptional programs, and TCIRG1-associated genetic variation converge on an autophagy-lysosomal module enriched in predicted non-responder subpopulations. The direct experimental evidence linking LAPTM5 to cisplatin resistance in NSCLC [70] further supports the functional relevance of this pathway in the PharmITH regulatory landscape.

#### Actin cytoskeleton pathway

The hub gene HCLS1 is aberrantly expressed in a drug non-responder LUAD subpopulation, suggesting activation of an actin-cytoskeleton remodeling program. HCLS1, normally an actin-binding signaling protein in immune cells, is upregulated in several cancers and has been linked to increased cell migration, invasion, and tumor formation [71]. In our GRN data, HCLS1 in non-responder cells is co-expressed with motility-, adhesion-, and immune-signaling genes, indicating a cytoskeletal and EMT-like transcriptional state.

Actin-network proteins are closely associated with malignant-cell motility, plasticity, and treatment-response phenotypes [72]. In our data, alterations in adhesion-related genes and splicing changes involving cytoskeleton-linked signaling provide additional regulatory features mapping to this pathway.

Together, HCLS1-associated transcriptional patterns and related post-transcriptional features converge on an actin-cytoskeleton remodeling module enriched in predicted non-responder LUAD subpopulations. These results position cytoskeletal plasticity as one of the recurrent pathway contexts associated with PharmITH, while HCLS1 serves as a hub connecting this pathway to the broader non-responder regulatory landscape.

### Supplementary Figures

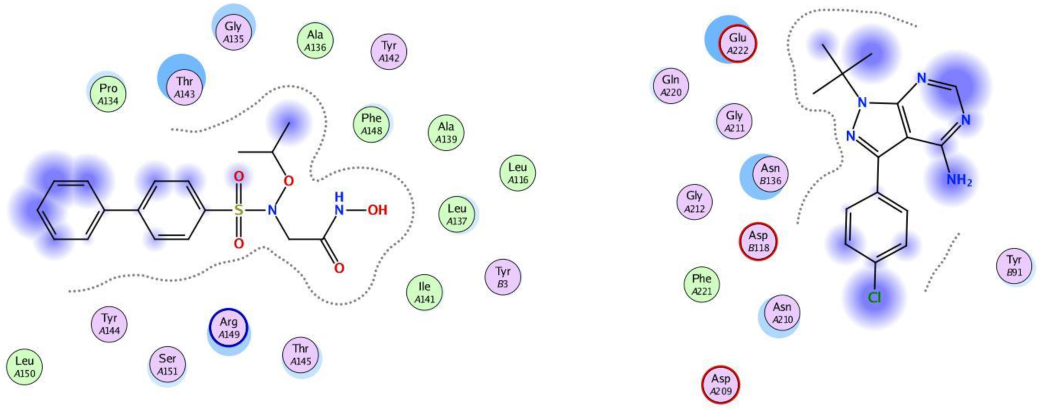


MMP2 Score: -6.8899 FYN Score: -5.3324

**Figure S1: Molecular docking of ARP-100 with MMP2 and PP2 with FYN.** The two-dimensional structures of ARP-100 and PP2 were downloaded from PubChem, saved in SDF format, and imported into MOE v2019 for energy minimization. The crystal structures of human MMP2 and FYN were obtained from the PDB database using Homo sapiens and a resolution of < 3 Å as the selection criteria. The protein structures were prepared by removing water molecules and other small molecules, and the inhibitors were docked with their corresponding targets in full-atom mode. The docking scores, calculated from the docking energies, indicate that ARP-100 and PP2 exhibit high binding affinity for MMP2 and FYN, respectively.

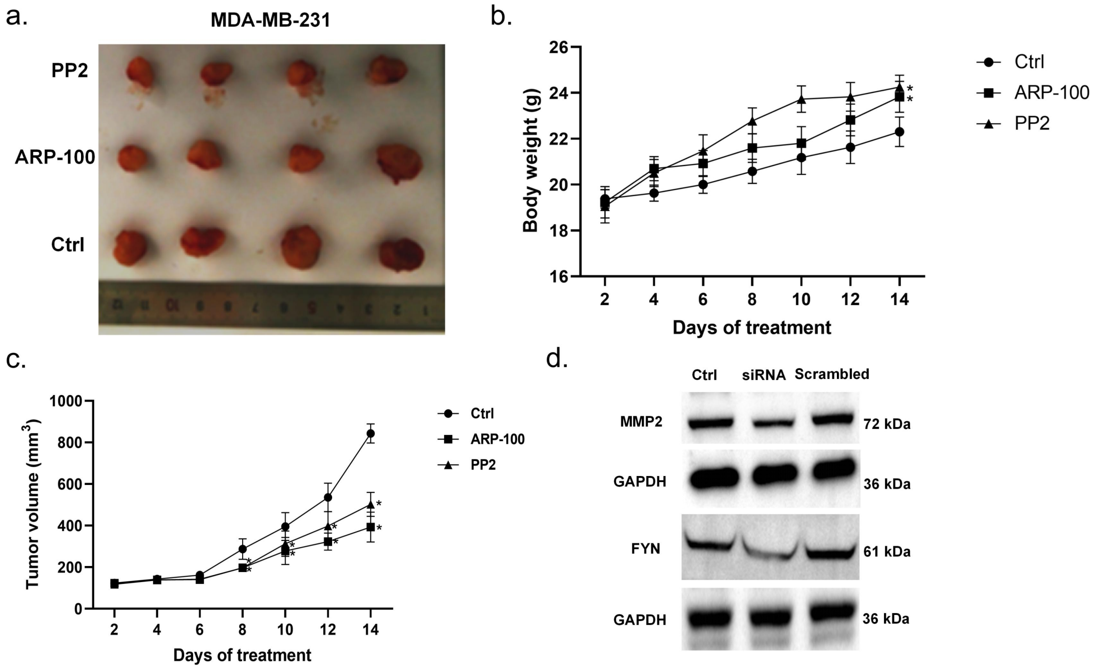


**Figure S2: Tumor growth and body weights of triple-negative breast cancer (TNBC) model.** **a.** Images of the tumor tissues statistically calculated in Fig 2.g. It can be observed that the tumors after inhibitor treatment show significantly slower growth compared to the original state. **b.** Mouse weights after the treatment, indicating the therapeutic effects of the inhibitors in subcutaneous tumor models. **c.** Tumor growth curve estimated from the tumor volumes. Results are mean ± SD (n = 4). * $p < 0.05$ compared with Ctrl group. **d.** Western blot analysis of MMP2 and FYN protein expression in the Ctrl, siRNA, and Scrambled groups. Ctrl represents the mock/no-siRNA control, and Scrambled represents the non-targeting siRNA control. GAPDH was used as the loading control.

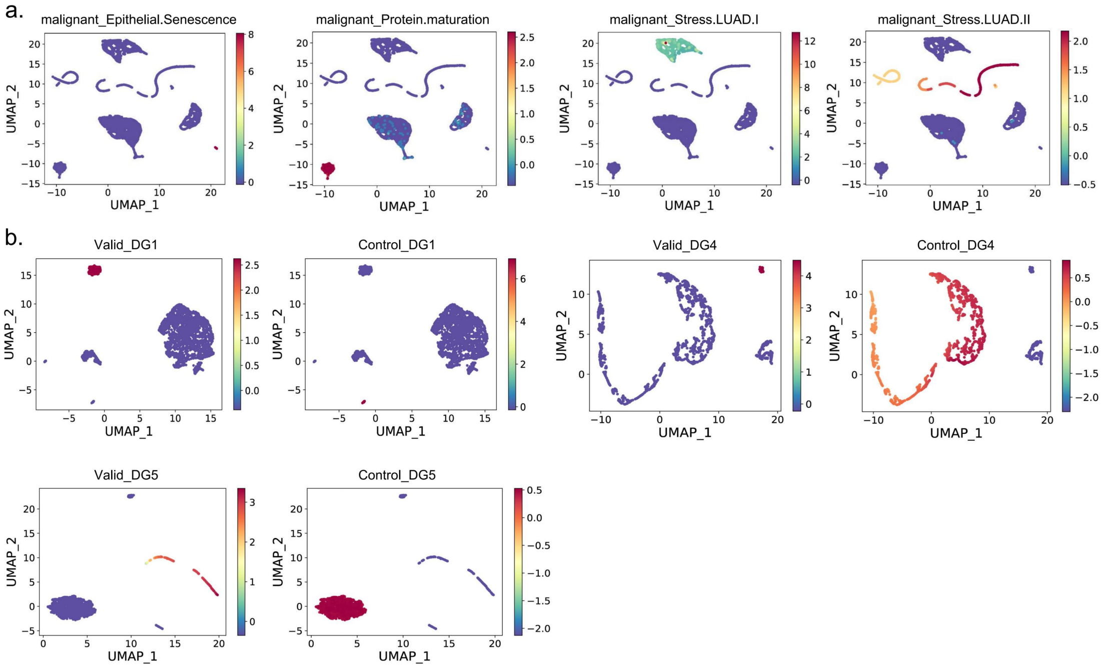


**Figure S3: Mapping DARs of Regulatory Heterogeneity within Tumors.** Defining differentially accessible regions (DARs) along two orthogonal dimensions: (a) at the ITH level, DARs that distinguish malignant meta-programs and capture intratumoral heterogeneity in regulatory potential; (b) at the PharmITH level, DARs that differentiate drug-sensitive from drug-resistant subpopulations within each drug group, thereby reflecting shifts in drug response. Results show significant differences in subtype-specific accessible regions across subpopulations, indicating that DARs contain clear PharmITH-driven factors.

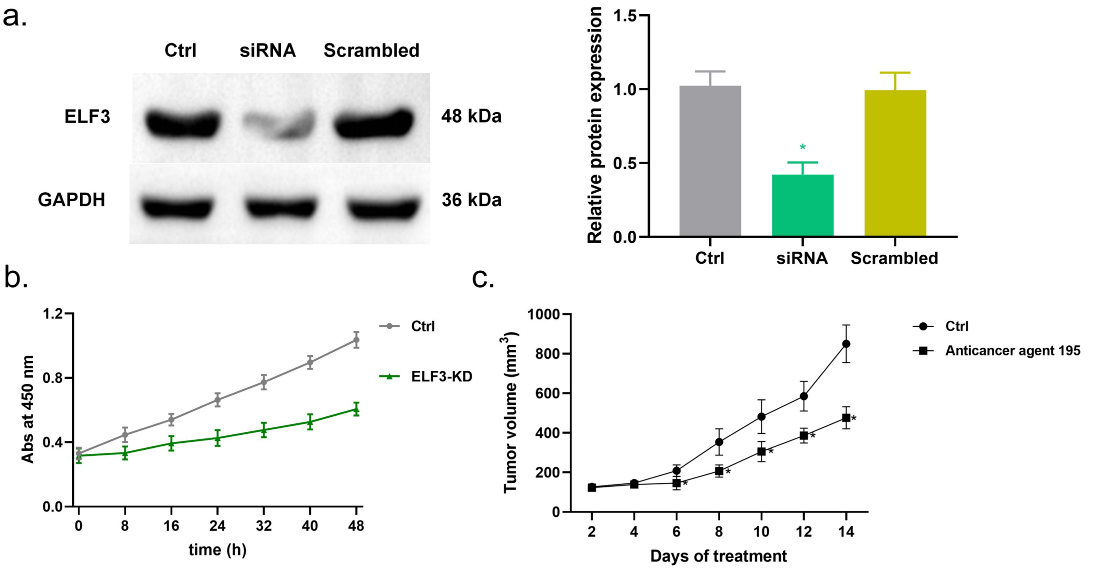


**Figure S4: Knockdown of ELF3 can significantly suppress tumor cell growth in LUAD cell line (A549).** a. Western blot and relative protein-expression analysis of ELF3 in A549 cells after ELF3 knockdown. Ctrl is the mock/no-siRNA control, and Scrambled is the non-targeting siRNA control. The bar graph shows the semi-quantitative relative ELF3 protein expression determined by ImageJ densitometric analysis. b. CCK-8 assay showing the proliferation and viability of A549 cells following ELF3 knockdown. Cell viability was evaluated by measuring absorbance at 450 nm over 48 h, with the time of medium replacement defined as time 0. c. Tumor growth curve estimated from the tumor volumes. Results are mean ± SD (n = 3 for in vitro experiments and n = 4 for animal studies). * $p < 0.05$ compared with Ctrl group.

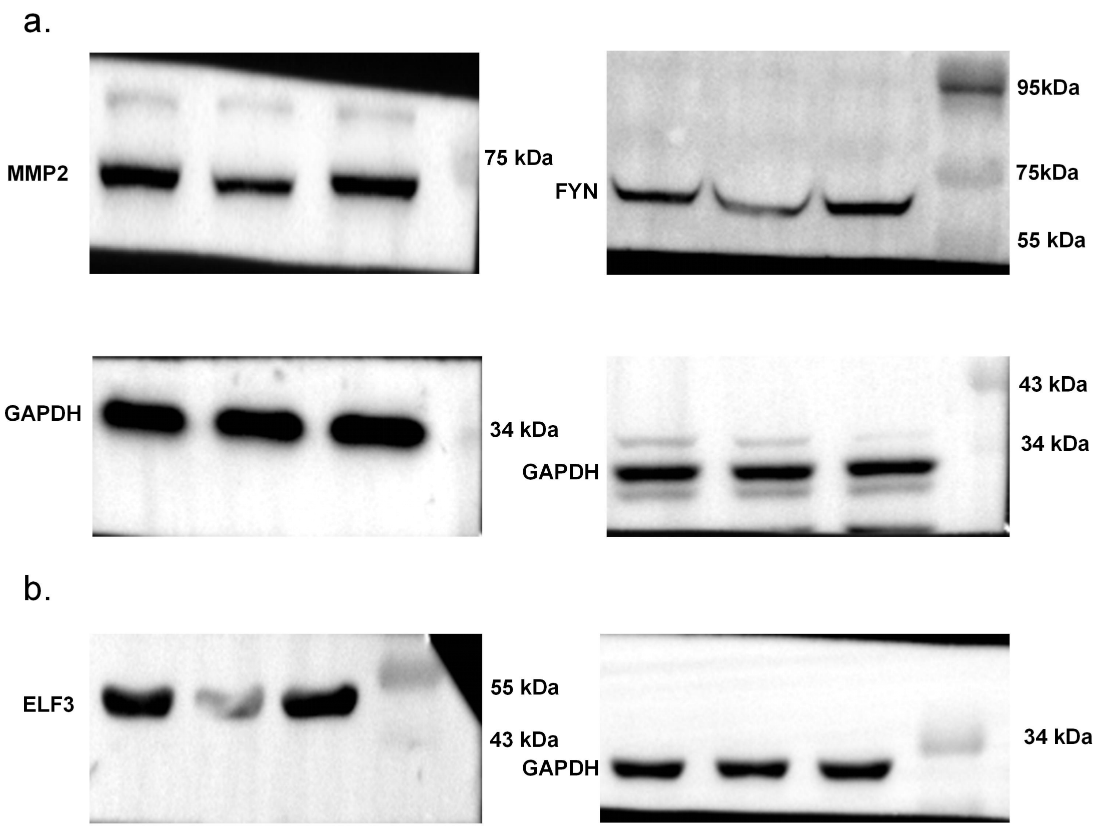


**Figure S5: Original western blot images with molecular marker bands.** a. Uncropped original blots for MMP2 and FYN in MDA-MB-231 cells. b. Uncropped original blots for ELF3 in A549 cells. Ctrl represents the mock/no-siRNA control, and Scrambled represents the non-targeting siRNA control. GAPDH was used as the loading control. Molecular weight marker bands are shown for each blot.

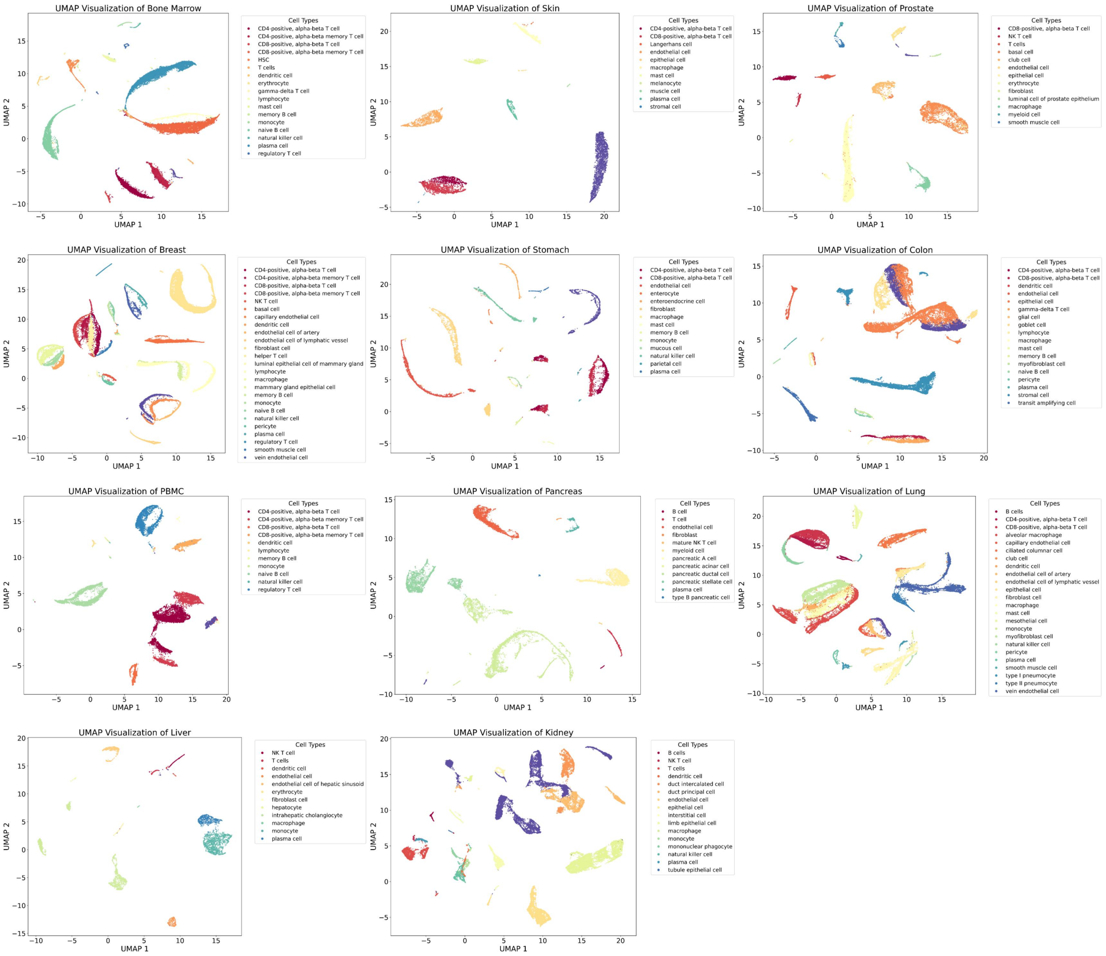


**Figure S6: Overview of cell types from scBERT.** To account for tissue specificity, we fine-tune our model using publicly available datasets derived from 11 distinct organs. All datasets are harmonized with respect to sequencing platform and other technical variables, and cell type labels are rigorously validated based on expert knowledge or experimental evidence. This approach ultimately enables the identification of 64 distinct cell types, encompassing major classes such as Immune cells, Myeloid, Erythroid, Stem cell, Epithelial, Endothelial, Mesenchymal, Neural and Tissue-specific populations.

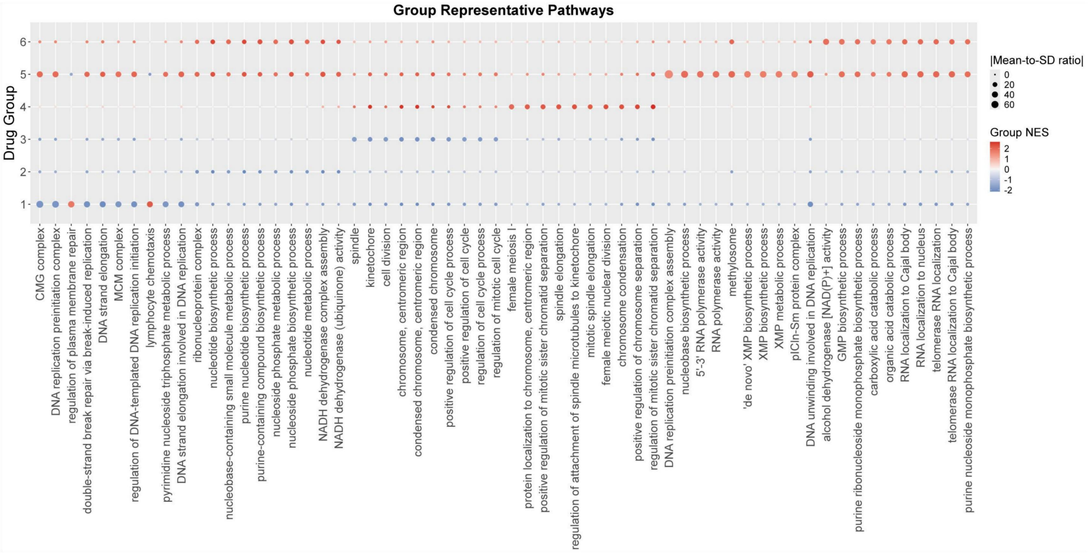


FigureS6.1 Drug group representative pathways of Acute Lymphoblastic Leukemia

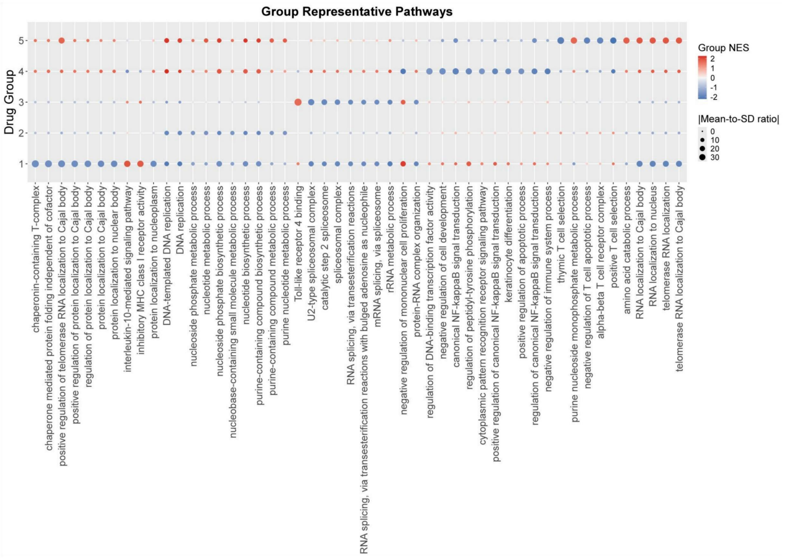


FigureS6.2 Drug group representative pathways of Acute Myeloid Leukemia-bone marrow

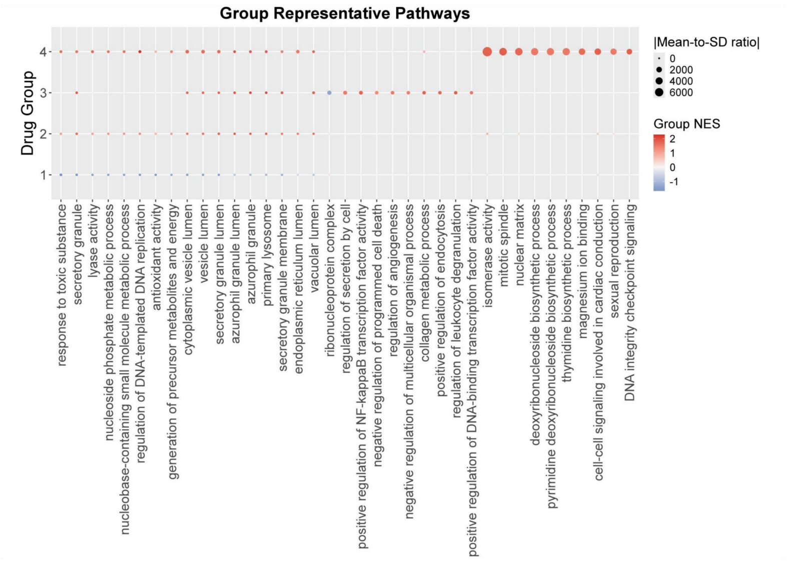


FigureS6.3 Drug group representative pathways of Acute Myeloid Leukemia-PB

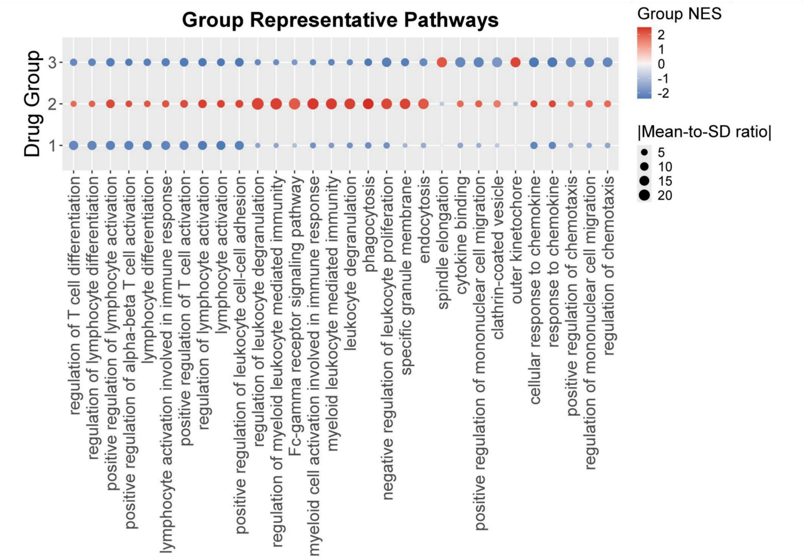


FigureS6.4 Drug group representative pathways of Breast Cancer

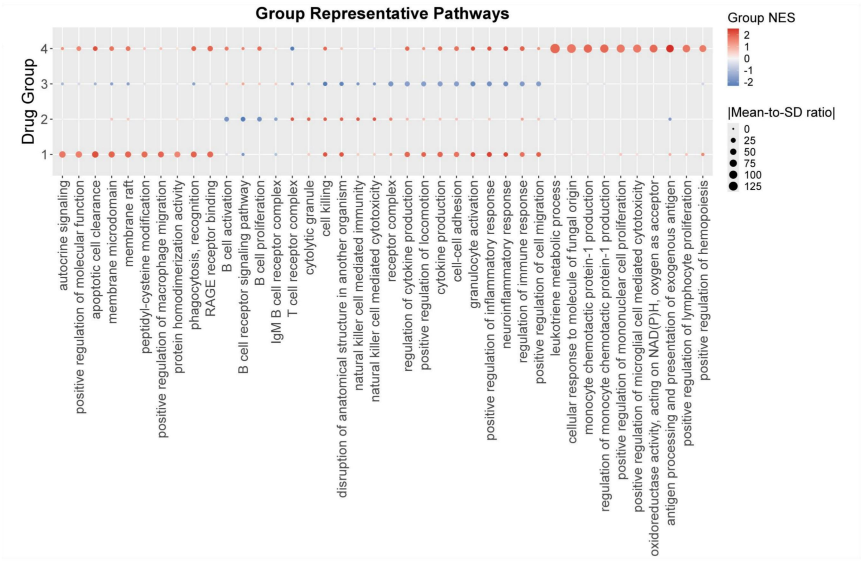


FigureS6.5 Drug group representative pathways of Chronic Lymphoblastic Leukemia

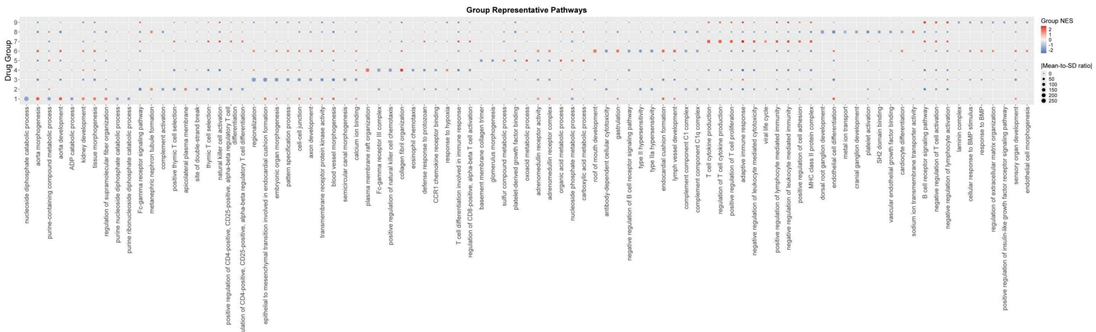


FigureS6.6 Drug group representative pathways of Clear Cell Renal Cell Carcinoma

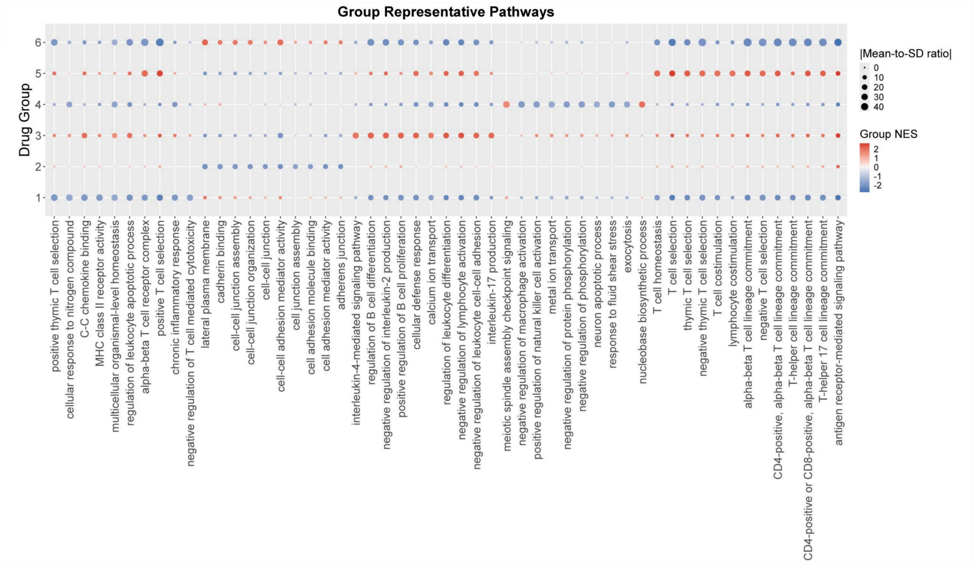


FigureS6.7 Drug group representative pathways of Colorectal Cancer

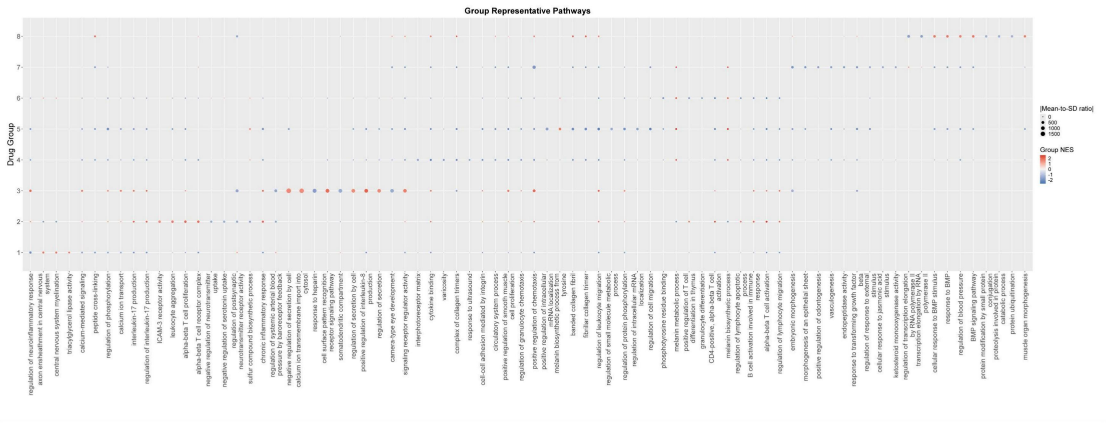


FigureS6.8 Drug group representative pathways of Cutaneous Melanoma

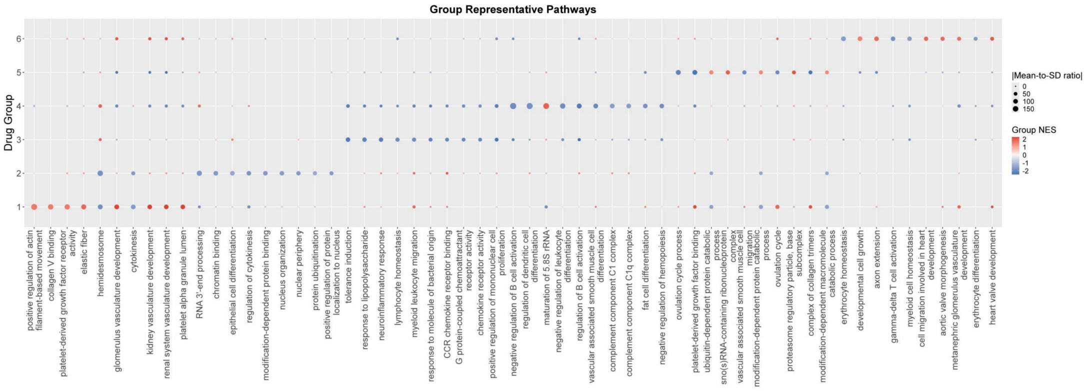


FigureS6.9 Drug group representative pathways of Cutaneous Squamous Cell Carcinoma

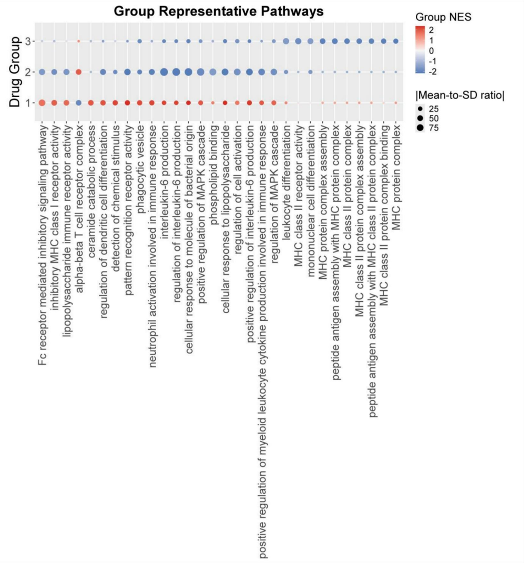


FigureS6.10 Drug group representative pathways of Cutaneous T Cell Lymphoma-PB

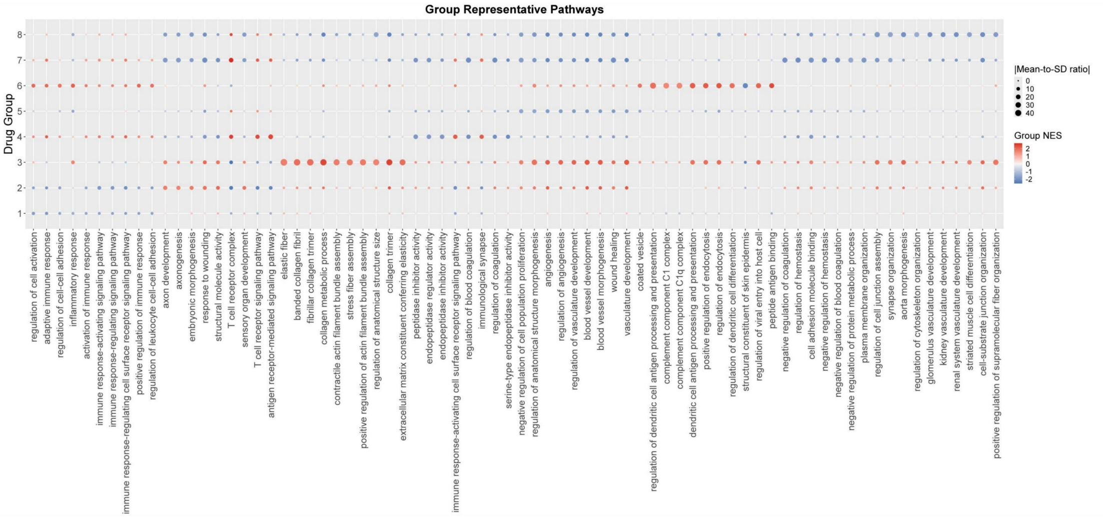


FigureS6.11 Drug group representative pathways of Cutaneous T Cell Lymphoma-Skin

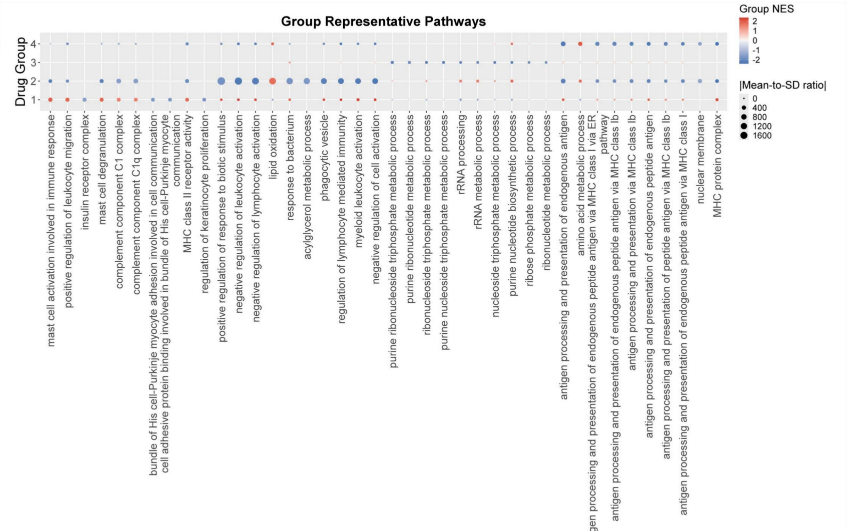


FigureS6.12 Drug group representative pathways of Ductal Carcinoma in Situ

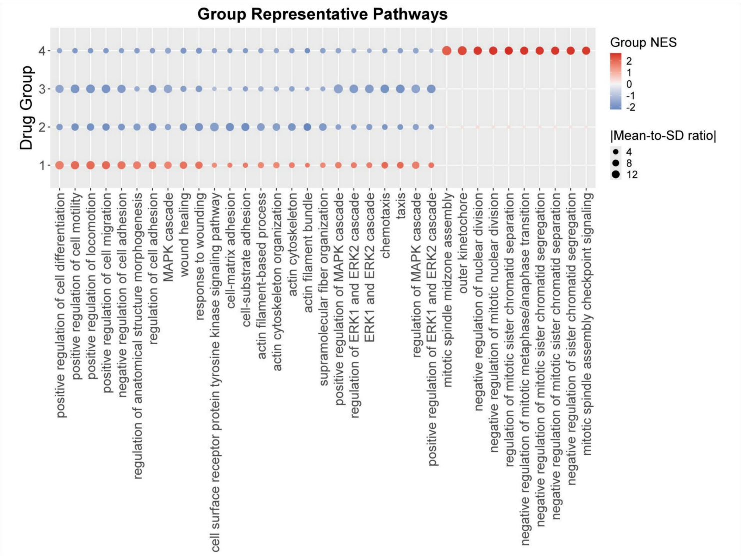


FigureS6.13 Drug group representative pathways of ER+ Breast Cancer

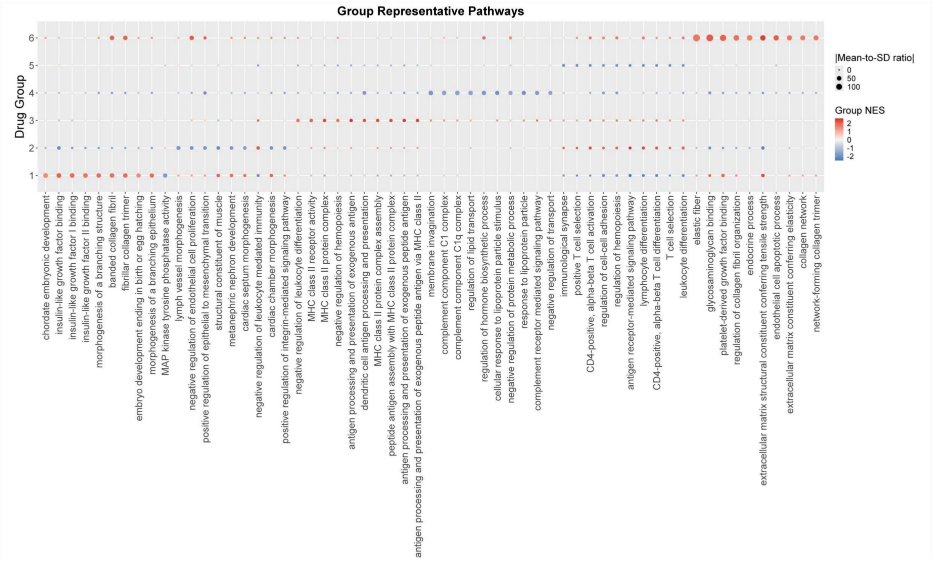


FigureS6.14 Drug group representative pathways of Gastric Cancer

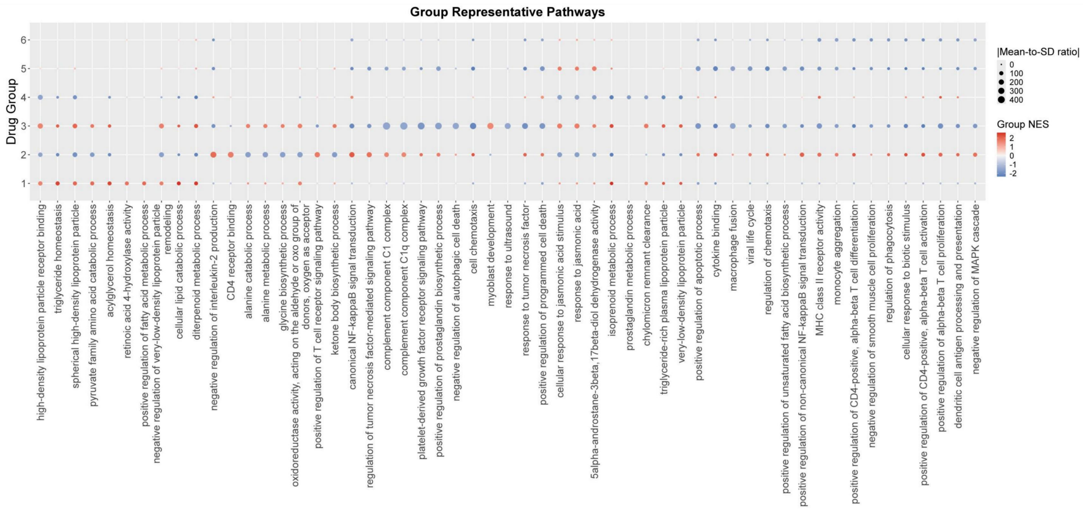


FigureS6.15 Drug group representative pathways of Hepatocellular Carcinoma

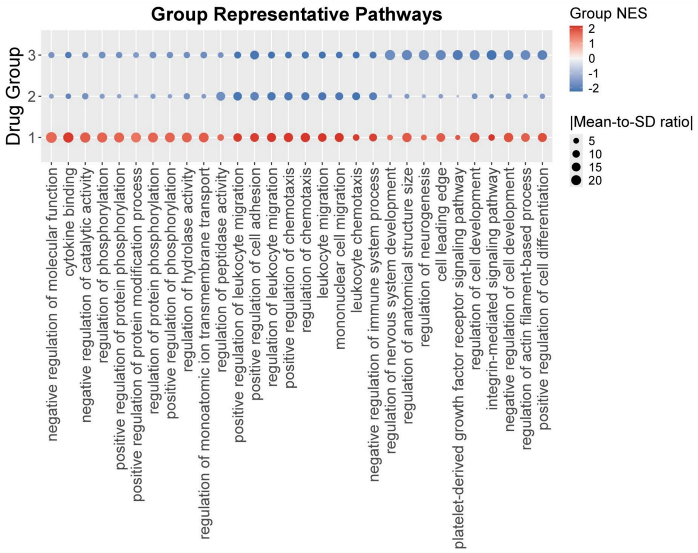


FigureS6.16 Drug group representative pathways of HER2+ Breast Cancer

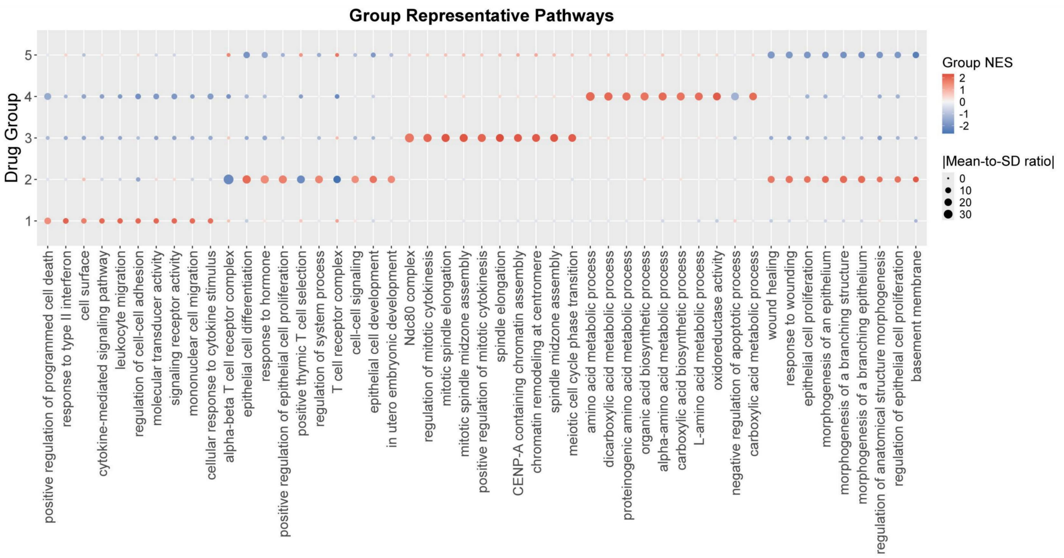


FigureS6.17 Drug group representative pathways of Intrahepatic Cholangiocarcinoma

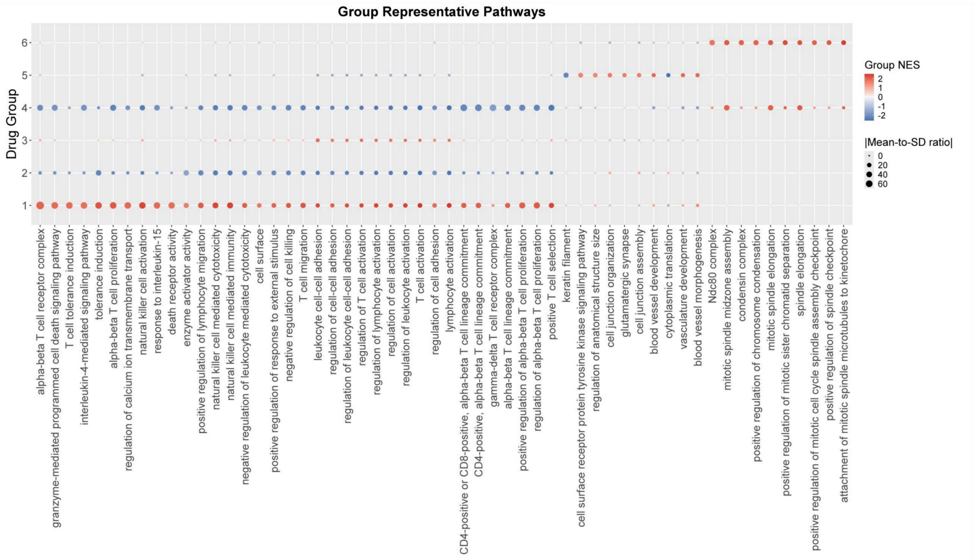


FigureS6.18 Drug group representative pathways of Invasive Ductal Carcinoma

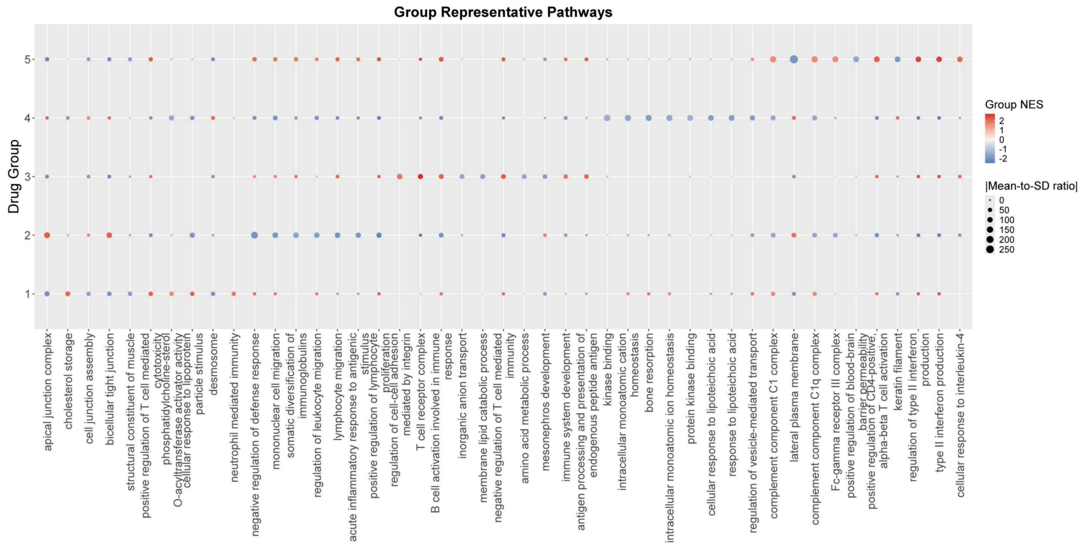

FigureS6.19 Drug group representative pathways of Lung Adenocarcinoma

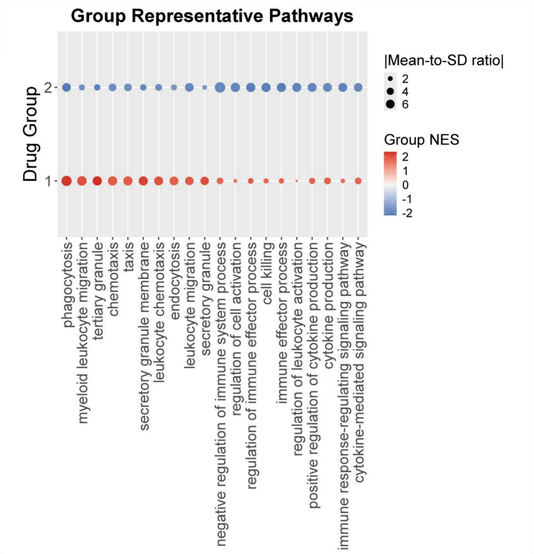

FigureS6.20 Drug group representative pathways of Lung Cancer

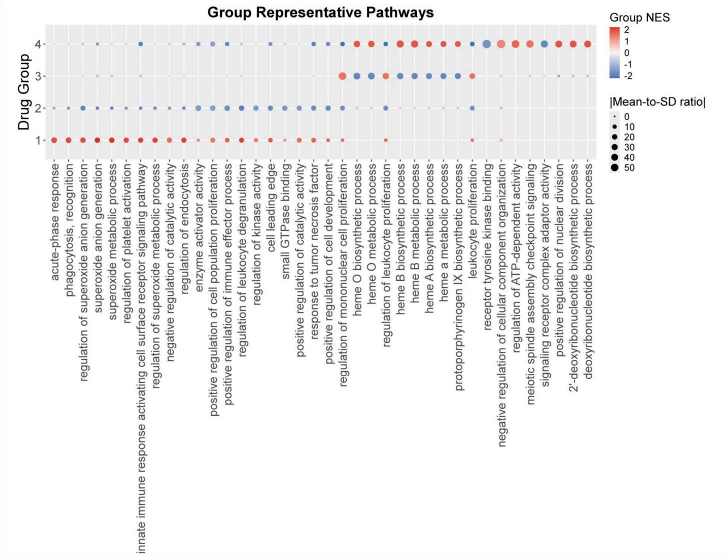


FigureS6.21 Drug group representative pathways of Myeloma-bone marrow

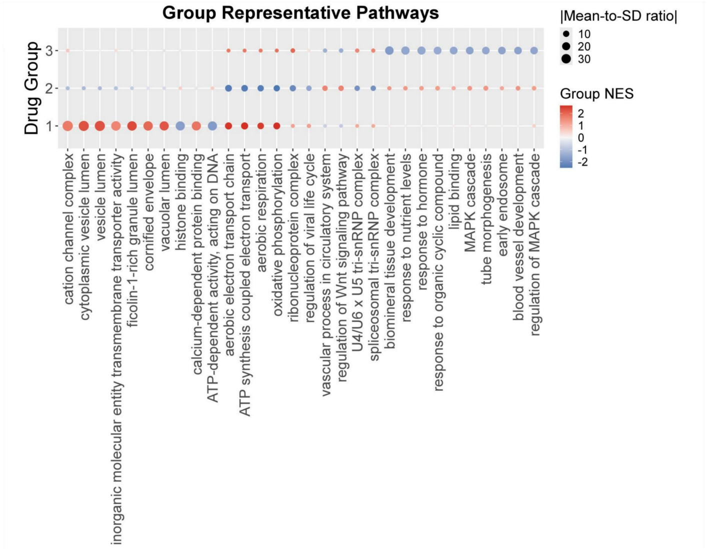


FigureS6.22 Drug group representative pathways of Myeloma-PB

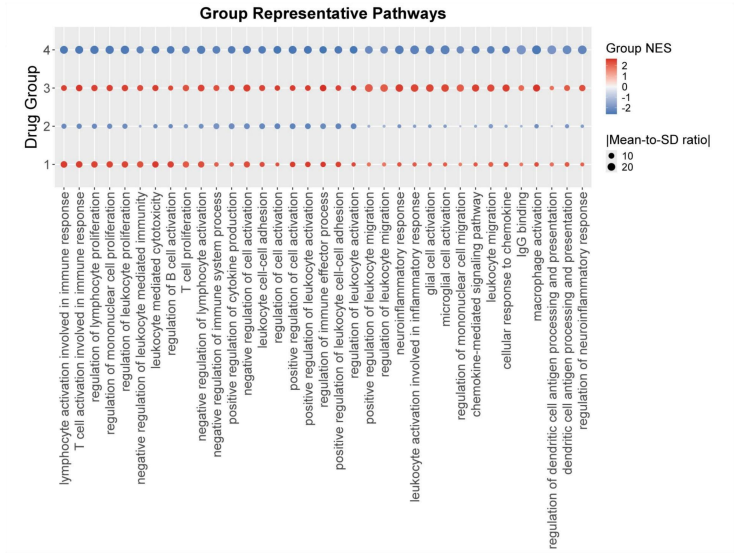


FigureS6.23 Drug group representative pathways of Non-Small Cell Lung Cancer

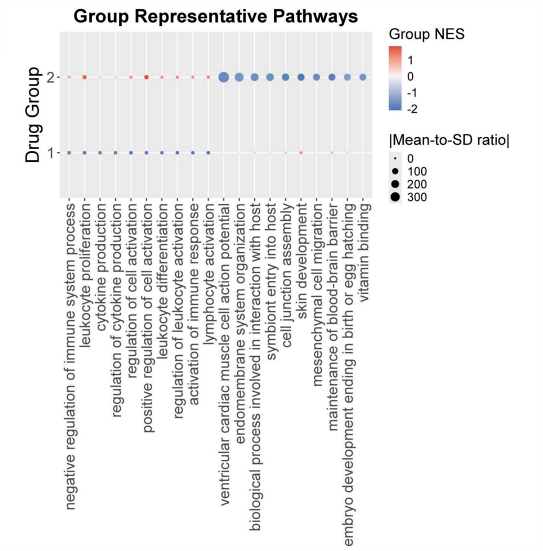


FigureS6.24 Drug group representative pathways of Pancreatic Cancer

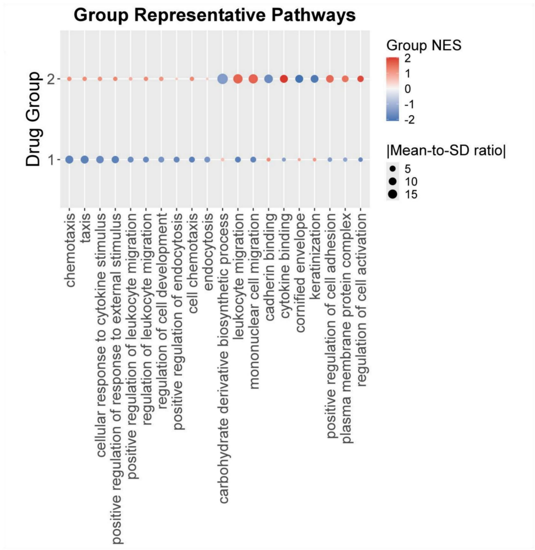


FigureS6.25 Drug group representative pathways of Pancreatic Ductal Adenocarcinoma

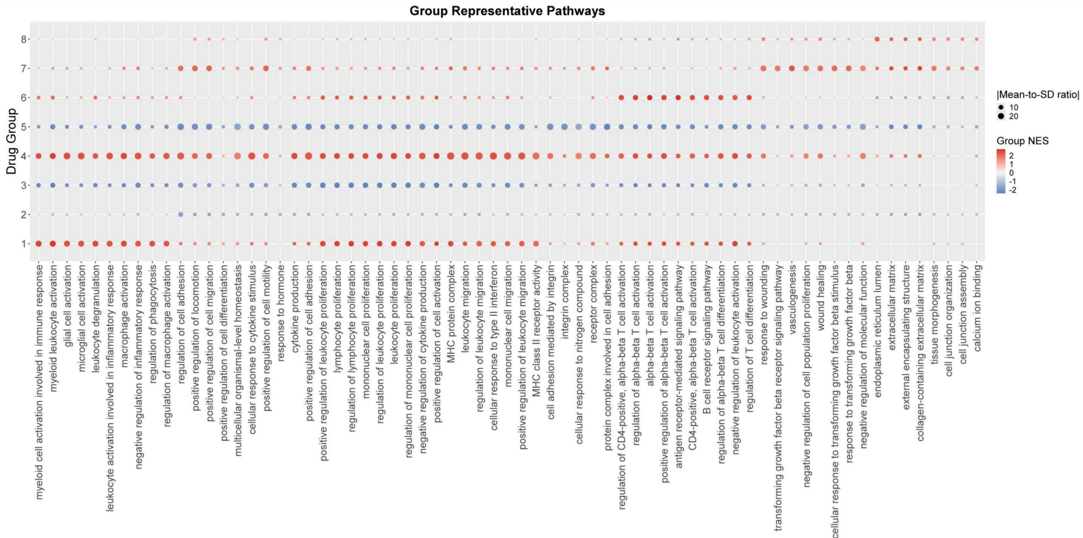


FigureS6.26 Drug group representative pathways of Prostate Cancer

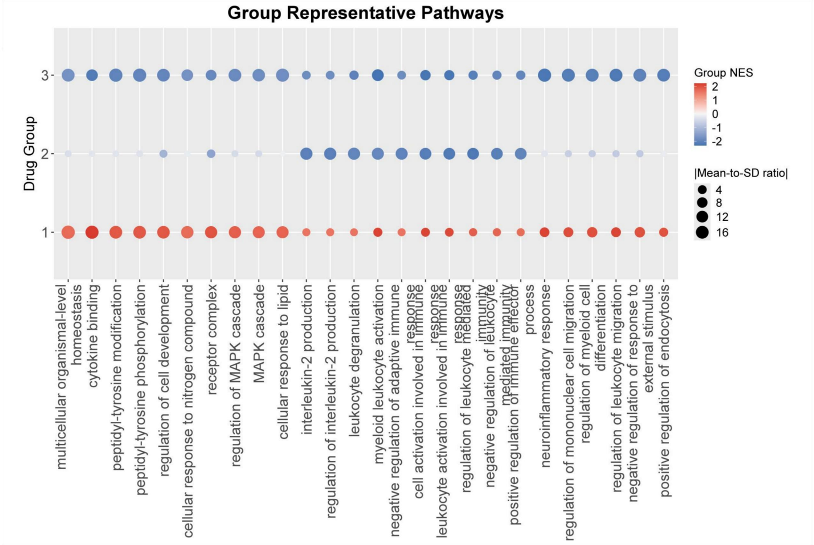


FigureS6.27 Drug group representative pathways of Triple-Negative Breast Cancer

**Figure S7: Overview of pathway enrichment across cancer drug groups.** GO pathways are used to enrich heterogeneous genes for each drug. Hierarchical clustering based on normalized enrichment scores (NES) groups drugs with similar mechanisms. (Due to extended size of this figure, we provide this figure as a separate PDF file)

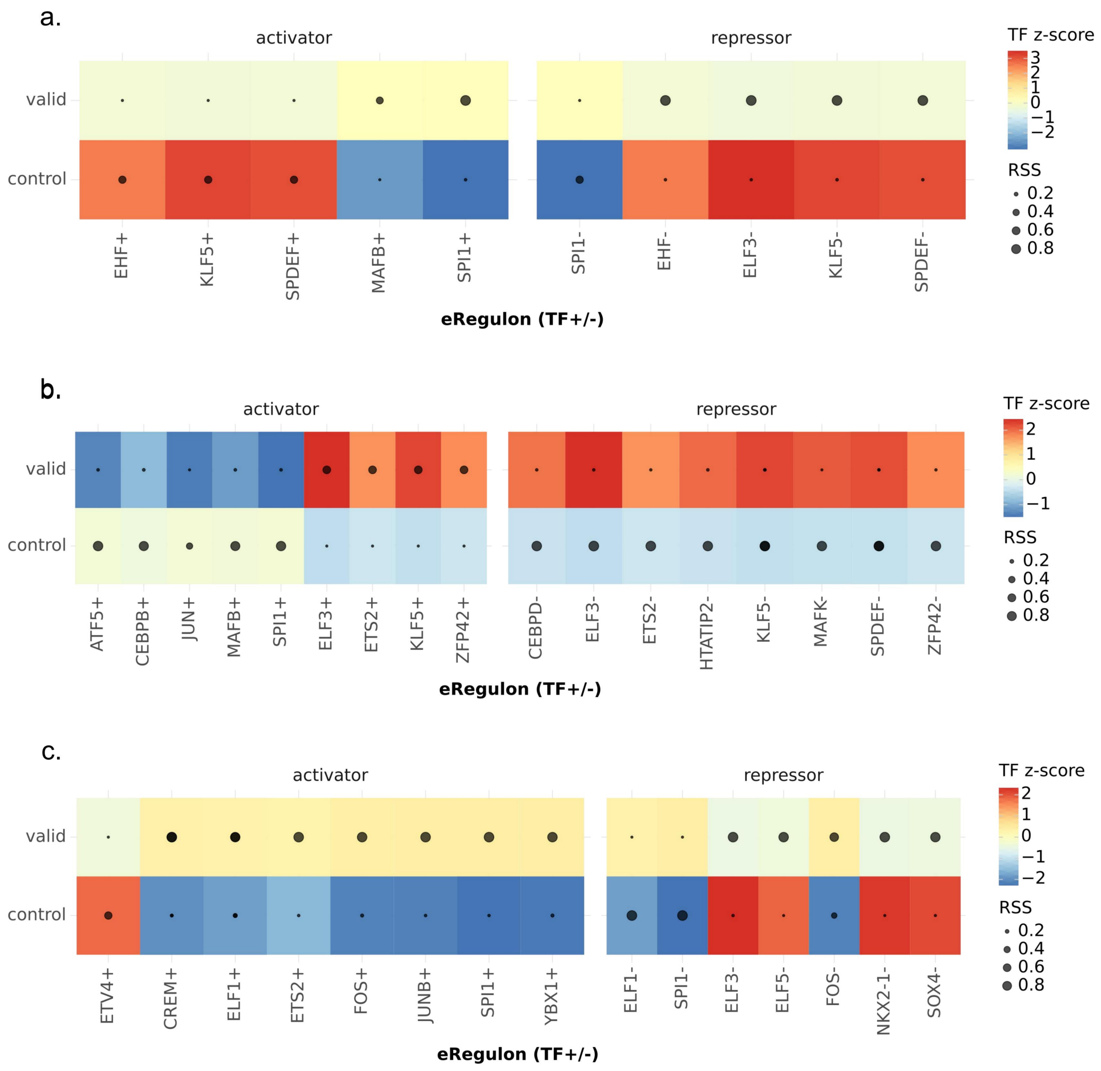


**Figure S8: Deciphering the transcription factor-hub gene regulatory network in PharmITH subpopulations.** a, b, and c represent drug groups DG1, DG4, and DG5, respectively. The symbols for TFs, '+' and '-', denote positive (activator) and negative (repressor) correlations with the hub gene. 'RSS' indicates the association between the hub gene and the corresponding cell subpopulation; 'TF z_score' is the normalized TF expression level.

### Supplementary Tables

Due to extended size of tables, we provide all tables as separate XLSX files. The caption of each table is as follows:

**Supplementary Table 1.** Overview of the 112 consensus malignant MPs identified across 24 major cancer types. The 'Programs' column details the functional annotations associated with each MP. Functional gene sets demonstrate MP-related specific gene sets.

**Supplementary Table 2.** List of the top 20 hub genes for each drug group in across 24 cancer types.

**Supplementary Table 3.** Literature survey (PubMed, 2018-2025) summarizing the experimentally validated functions of selected hub genes in specific cancer contexts. 'Direct Therapeutic Target' indicates genes validated as therapeutic targets; 'Biomarker' indicates genes serving as predictive, prognostic, or diagnostic markers; 'Immune Process' indicates genes directly involved in immune cell behavior, while chemokines and antigen presentation genes annotated as tumor-associated functions.

**Supplementary Table 4.** Top 100 high-confidence Transcription factor (TF) – Enhancer – Target hub gene trios inferred by SCENIC+ from the GSE274934 project (paired scRNA-seq and scATAC-seq). In the 'Transcription factor' column, '+' and '-' denote positive and negative correlation with the hub gene, respectively. 'gene_based_RSS' indicates the association between the hub gene and the corresponding cell subpopulation; 'TF_expression_zscore' is the normalized TF expression level; 'Enhancer region' denotes the differentially accessible enhancer region linked to the hub gene. Analyses were performed separately for ITH and PharmITH subpopulations.

**Supplementary Table 5.** List of 41 high-significance ($p\_val < 0.05$) somatic mutation events associated with PharmITH from the GSE274934 project. 'Event' specifies the genomic coordinate and mutation; 'Category' denotes the associated subpopulation (all linked to the drug-responder group in this analysis); 'Func.refGene' provides the functional category of the mutated region; 'GeneDetail.refGene' gives the distance to the nearest gene for upstream/downstream/intergenic mutations; 'ExonicFunc.refGene' details the functional consequence for exonic mutations; 'AAChange.refGene' provides transcript-specific amino acid change information.

**Supplementary Table 6.** Hub gene-mediated cell-cell communication events in the GSE274934 project. Analyzed across three contexts: general malignant ITH, drug-responder, and drug non-responder subpopulations interacting with the tumor microenvironment.

**Supplementary Table 7.** Significant alternative splicing events associated with drug sensitivity groups in the GSE274934 project. 'Drug sensitivity' labels the subpopulation ('control' for non-responder, 'valid' for responder). 'ScZ_median' quantifies the deviation of splicing patterns in the subpopulation compared to the entire cell population. Positive values indicate the splicing preference for longer introns, while negative values indicate the preference for shorter introns.

**Supplementary Table 8.** Hub gene-associated regulatory interactions inferred using SCENIC across 24 cancer types. 'importance' quantifies the contribution of the transcription factor (TF) to the target hub gene's expression; 'imp_pct_tf' represents the percentile rank of the importance value for the target hub gene among all candidate targets of the same TF; 'active_frac' indicates the fraction of cells within a population where the TF regulon is deemed active.

**Supplementary Table 9.** Cell-cell communication events where hub genes function as ligands or receptors between malignant and tumor microenvironment cell populations. 'specificity_rank' ranks the significance of the ligand-receptor pair (lower value indicates higher specificity); 'magnitude_rank' ranks the communication probability strength of the pair (lower value indicates greater strength).

**Supplementary Table 10.** Cell-cell communication events mediated by hub genes as ligands or receptors, specifically from drug-responder ('_valid') or drug non-responder ('_control') malignant subpopulations to the tumor microenvironment.

ICAM-1 protects lung carcinoma from apoptosis through ligation of fibrinogen. Cell Death & Disease, 15(8), 605.

63. Qiang, M., Chen, Z., Liu, H., Dong, J., Gong, K., Zhang, X., ... & Tang, M. (2025). Targeting the PI3K/AKT/mTOR pathway in lung cancer: mechanisms and therapeutic targeting. Frontiers in pharmacology, 16, 1516583.
64. Sanaei, M. J., Razi, S., Pourbagheri-Sigaroodi, A., & Bashash, D. (2022). The PI3K/Akt/mTOR pathway in lung cancer; oncogenic alterations, therapeutic opportunities, challenges, and a glance at the application of nanoparticles. Translational Oncology, 18, 101364.
65. Cyran, A. M., & Zhitkovich, A. (2022). Heat shock proteins and HSF1 in cancer. Frontiers in oncology, 12, 860320.
66. Mao, H., Zhao, X. & Sun, Sc. NF-κB in inflammation and cancer. Cell Mol Immunol 22, 811-839 (2025). https://doi.org/10.1038/s41423-025-01310-w
67. Wang, H., Xiong, A., Chen, X., Guo, J., Tang, Z., Wu, C., ... & Jiang, T. (2024). CXCR1+ neutrophil infiltration orchestrates response to third-generation EGFR-TKI in EGFR mutant non-small-cell lung cancer. Signal transduction and targeted therapy, 9(1), 342.
68. Ibusuki, R., Iwama, E., Shimauchi, A., Tsutsumi, H., Yoneshima, Y., Tanaka, K., & Okamoto, I. (2024). TP53 gain-of-function mutations promote osimertinib resistance via TNF-α-NF-κB signaling in EGFR-mutated lung cancer. NPJ precision oncology, 8(1), 60.
69. Wu, D. D., Dai, L. J., Tan, H. W., Zhao, X. Y., Wei, Q. Y., Zhong, Q. H., ... & Xu, Y. M. (2022). Transcriptional upregulation of MAPK15 by NF-κB signaling boosts the efficacy of combination therapy with cisplatin and TNF-α. Iscience, 25(12).
70. Wu, F., Li, C., Song, X., & Xie, L. (2025). LAPTM5 confers cisplatin resistance in NSCLC by suppressing LAMP1 ubiquitination to stabilize lysosomal membranes and sustain autophagic flux. Cellular Signalling, 111834.
71. Koya, Y., Liu, W., Yamakita, Y., Senga, T., Shibata, K., Yamashita, M., ... & Kajiyama, H. (2018). Hematopoietic lineage cell-specific protein 1 (HS1), a hidden player in migration, invasion, and tumor formation, is over-expressed in ovarian carcinoma cells. Oncotarget, 9(66), 32609.
72. Shen, L. P., & Jiang, H. T. (2024). Pan-cancer and single-cell analysis of actin cytoskeleton genes related to disulfidptosis. Open Medicine, 19(1), 20240929.
